# Inducing highly physiologically relevant phenotypes of human vascular smooth muscle cells via 3D printing

**DOI:** 10.1101/2020.07.24.206888

**Authors:** Peiran Zhu, Xuzhao Li, Wang Xin, Menglin Wang, Chengzhen Yin, Jinze Li, Hangyu Chen, Hengjia Zhu, Yubing Sun, Jiemin Jia, Nanjia Zhou

## Abstract

Vascular smooth muscle cells (vSMCs) are one of the essential cell types in blood vessel walls. A significant vSMC phenotype characteristic is that they collectively wrap around the outer layer of the healthy blood vessels with spindle-like morphology and help maintain the vascular tones and regulate the blood flow. Both physiological and biomedical research are impeded by the standard 2D cell culture approaches which do not create *in vivo* like microenvironment. Here, we systematically investigated the vSMCs culturing within 3D printed geometrical constraints and on printed microfilaments. Based on these models, we demonstrate a simple bioprinting approach for fast manufacturing vessel architectures with micro-grooved surfaces for vSMCs alignment. We validated that the vSMCs cultured on the printed vessel with microfilaments (VWMF) present a more physiologically relevant morphological phenotype and gene expression profile, and they are considerably more active in wound healing and ischemia than conventional planarly cultured vSMCs.

## INTRODUCTION

In blood vessels of mammals, vSMCs are highly aligned circumferentially along the vessels and they adopt a quiescent, contractile phenotype to maintain vascular tone and regulate blood flow^1^ (**Figure 1a-b**). Specifically for vSMCs, upon rupture of vessels^2^ or during ischemia^5^, the microenvironmental changes may result in a transition from a contractile phenotype to a synthetic migratory phenotype, which eventually leads to the development of cardiovascular diseases (CVD)^3–5^. Microenvironment mediated cellular phenotype transition has been investigated using conventional in vitro cell culture^6–11^. However, standard cell culture models do not create 3D microenvironment^12^. Lacking the relevant biomanufacturing methodologies, the phenotype regulatory mechanisms for inducing *in vivo*-like vSMCs *in vitro* are still unclear.

**Figure 1.**
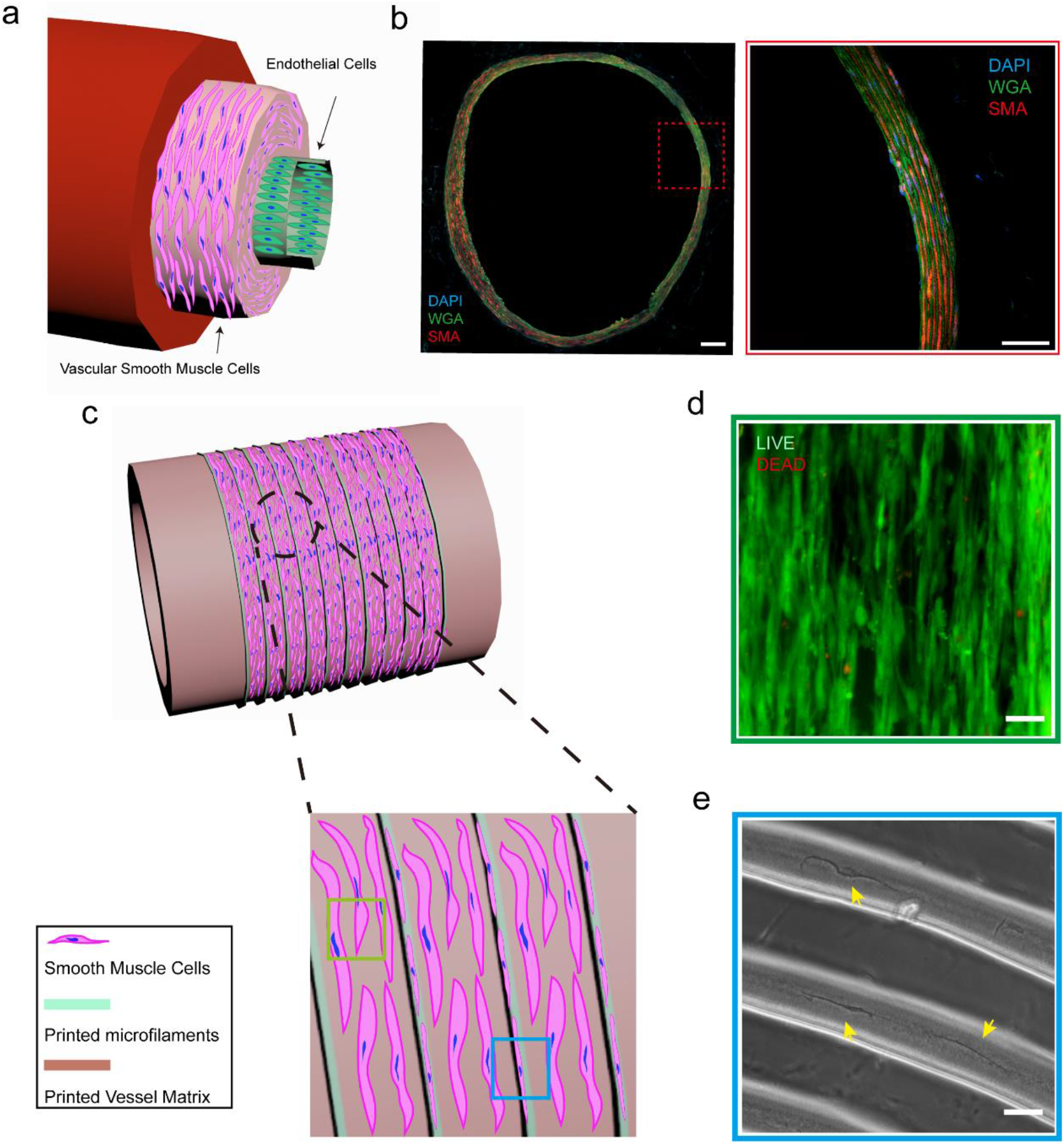
Inducing biomimetic smooth muscle layer through 3D culturing of vSMCs on VWMFs. (a) Schematic diagram showing the real structure of mammal aorta blood vessel, which mainly contains three layers, including vascular endothelial cells, vascular smooth cells, and tunica externa. (b) Representative fluorescence images showing a slice of *NG2DsRed* mouse aorta vessel. Scale bar, 100 μm (left) and 50 μm (right). (c) Schematic diagram presenting the 3D printed VWMF. The enlarged figure indicates that vSMCs are highly aligned to the orientation of the printed filaments. (d) Representative fluorescence image showing the highly oriented vSMCs cultured on VWMFs with high viability through staining with LIVE/DEAD reagent. Scale bar, 50 μm. (e) Representative phase contrast image showing the vSMCs expended along the top of printed microfilaments. Scale bar, 50 μm.

The demand for artificial blood vessel implantation for various types of cardiovascular disorder spurs significant interests in new 3D biomanufacturing methodologies. As the function of the vessels largely rely on the geometries of the host vasculatures for supporting functional cell culture, these methods must simultaneously provide the vessel support and microenvironment necessary for 3D cell culture. Recently, different manufacturing strategies such as micro-molding, electrospinning, and 3D printing have been investigated for *in vitro* vessel manufacturing^13–16^. Among them, 3D printing technologies provide the advantages of fabricating custom-shaped architectures using engineered materials, and they are highly ideal for manufacturing implantable blood vessels. For instance, Zhang et al.^17^ generated the coagulation models to investigate the pathology of fibrosis in thrombosis. Cui et al.^18^ printed a small-diameter vasculature through a core-shell nozzle, which including smooth muscle and endothelium cells and evaluated the performance both *in vitro* and *in vivo*. While these methods are successful in generating various 3D vessel-like geometries, they do not sufficiently recapitulate the vessel structures with circumferentially aligned vSMCs phenotypes *in vitro*, which seriously impeded the further investigation of physiological disease derives from the dysfunction of smooth muscle cells.

Here, to systematically investigate how microenvironment mediates the cellular phenotypes of vSMCs, both digital light processing (DLP) and direct ink writing (DIW) were employed to create various 2D and 3D ECM patterns. While the standard 2D cell culture methods do not induce spontaneous alignment of vSMCs (**Supplementary Figure 1**), we investigated the modulation of cell behavior by 2D/2.5D geometrical constraints. Specifically, we engineer the anisotropic vSMCs within printed rectangular-shaped wells and on printed microfilaments. Employing a new printhead design, we demonstrate a simple, one-step approach to print the complete biomimetic vessel architecture with external micro-grooved surface for vSMCs self-assembly. After coating human fibronectin on VWMF to facilitate cell adhesion, we successfully generated smooth muscle layers consisting of circumferentially oriented vSMCs on the external wall of the printed vessels (**Figure 1c-e**) and induced a highly contractile phenotype of vSMCs which show high cellular aspect ratio. The molecular studies further confirm that the 3D culturing on VWMF induce a more *in vivo* like vSMCs with a quiescent phenotype. We finally created pathological models, including wound healing assay to mimic vascular trauma and the standard oxygen glucose deprivation (OGD) assay to simulate the ischemic environment *in vivo*. Our 3D cultured vSMCs on VWMF exhibit enhanced proliferation after injury and a choppier response to ODG than 2D cultured vSMCs.

## RESULTS

### Microenvironment prominently modulates the phenotype and manners of vSMCs

We first employ DLP technology to print 3D stencils to confine vSMCs in different planar geometries (**Supplementary Figure 2**). We primordially cultured vSMCs on a circular pattern with a diameter of 400 μm. Different from the previous reports that various types of cells, including stem cells, cancer cells, endothelium cells, are found to distribute uniformly in the whole pattern^9, 12^, we found that vSMCs are predominantly crowded in the center of the pattern after two days. Increasing cell seeding density from 160 to 450 cells/mm^2^ (**Supplementary Figure 3**) shows the same trend. Not surprisingly, the inhomogeneous distribution of vSMCs disappeared with the increasing of pattern dimension (**Supplementary Figure 4**). Since cellular migration and expansion could be regulated by the cytoskeleton and intercellular tensions^19, 20^, we next treated cells with Y27632 (10 μM, dosed 1 h after cell seeding) to reduce cytoskeletal tensions and yielded a homogeneous distribution of vSMCs (**Supplementary Figure 5, Figure 2a**). In contrast to the cells crowded in the pattern center, the cells distributed in the periphery were highly expanded, and the nuclear, cytoskeleton and stress fibers all conform to the patterned curvature (**Supplementary Figure 6a**). Consistent with the previous results, this phenomenon disappears after ROCK inhibitor treatment (**Supplementary Figure 6b**).

**Figure 2.**
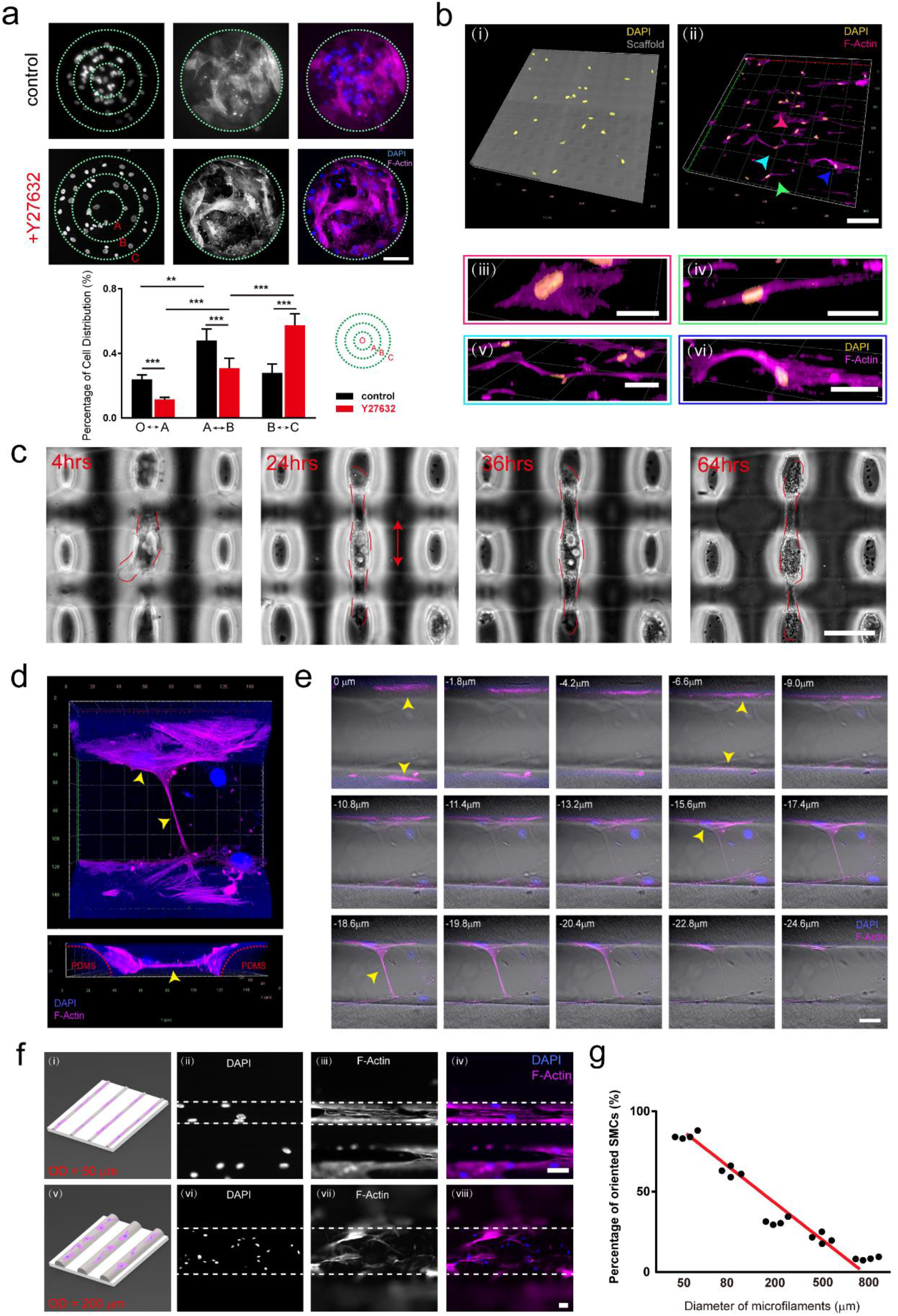
Microenvironment modulating the behaviors of vSMCs. (a) Representative fluorescence images (top) showing vSMCs cultured on a 2D fixed pattern. Cells were stained with phalloidin (pink) and DAPI (blue). Scale bar, 100 μm. Bar plot (below) showing the percentage of vSMCs distributed in the different region of the pattern. Data represent mean ± s.e.m. from at least three independent experiments. *, P < 0.05, **, P < 0.01, ***, P < 0.001, n.s., P > 0.05. (b) Representative confocal images showing cells on printed 3D scaffold for 3 days. Cells were stained with phalloidin (pink) and DAPI (yellow). Scale bar, 100 μm for (i-ii), 25 μm for (iii-iv), and 50 μm for (v-vi). (c) Representative phase contrast images showing the unidirectional elongation of vSMC through time-lapse imaging. Red dotted lines mark the outline of the cells. Scale bar, 100 μm. (d) Representative confocal images showing top view (top) and side view (below) of vSMC suspended between two microfilaments on the printed scaffold. Cells were stained with phalloidin (pink) and DAPI (blue). (e) Z-stack images for (d) obtained from surface to bottom of the scaffold (thickness = 2.4 μm). Scale bar, 20 μm. (f) Schematic diagrams (left) and representative fluorescence images (right) showing the vSMCs cultured on top of the printed filaments. Cells were stained with phalloidin (pink) and DAPI (blue). Scale bar, 50 μm. (g) Scatter plot showing the percentage of oriented vSMCs as a function of the diameter of the filaments, and the result of linear fitting.

To further investigate the effect of size and aspect ratio of controlled microenvironment on the cell culture, we next print more complex geometries. We printed planar grids with variable areas and aspect ratios, and found that vSMCs are able to fully occupy small grids with areas <1600 μm^2^ but are only able to cover the larger grids partially. When cells cultured in a rectangular grid with a high aspect ratio, cells expanded and elongated along the sides of the grid (**Supplementary Figure 7**). We conclude that the size and aspect ratio of restricting ECMs could direct the expansion and elongation of vSMCs. We next printed multilayer 3D scaffolds (**Supplementary Figure 8a-b**), and more cellular shapes were detected after culturing of vSMCs for 3 days. Through fluorescence confocal imaging (**Figure 2b i-ii**), we found more types of vSMCs, including homogeneously expanded cells, highly elongated cells, and even spindle-like cells stretching over and binding to several printed microfilaments with a high cellular aspect ratio similar to their behaviors *in vivo* (**Figure 2b iii-vi, Supplementary Figure 8c**). For these spindle-like cells, we captured the growth process of a sac by live-cell imaging. The results indicated that the cell first attached to the bottom of the scaffold and then vertically elongated in the following 64 hrs and eventually attached to two adjacent microfilaments, exhibiting a high cellular aspect ratio of 7 (**Figure 2c**). We further observed that the cells could form an extremely fine cell process that suspended between two adjacent microfilaments and left the cell body in the two sides (**Figure 2d-e**).

These results from patterned cell culture studies suggest that surface topologies such as curvatures of the 3D printed ECMs significantly influence the vSMC culturing. Recent studies reported that the ECM curvature could direct the ROCK signaling and migration of vSMCs^21^. We next directly printed microfilaments with different nozzle diameters from 50 μm to 800 μm on coverslips. The convex geometry resulted from semicircular cross-sections of the printed filaments delivered a decreasing curvature with the rising of filament diameter for vSMCs, and the results showed that a steeper geometry could direct the elongation and orientation (along the radial direction) of vSMCs. Furthermore, within the increasing of filament diameter, vSMCs expanded more disorderly and presented a smaller aspect ratio (**Figure 2f-g, Supplementary Figure 9**). Paradoxically, the vSMCs collectively wrap around the vascular wall in vivo, which means that it is impossible to realize circumferential orientation of vSMCs in vitro through simply providing ECM with tubular geometries. Additionally, we next explored how concave geometry directs the manners of vSMCs through printing fugitive material (Pluronic F127) onto the glass. After pouring PDMS and incubating two hours under 80 °C, the concave matrix could be achieved after washing off sacrificial material. Interestingly, the vSMCs spontaneously wrapped along the circumferential direction in a collective manner after 40 hrs (**Supplementary Figure 10**). Altogether, these results suggest that the curvature of ECM could dramatically mediate the vSMC phenotypes, especially the cellular aspect ratio, which was recognized to be highly correlated with the capacity of vSMCs contraction^6^.

### Collectively alignment of vSMCs through printing tailored microfilaments

Structure determines functions. Collectively circumferential alignment of vSMCs is regarded as the foundation of structural maintenance and spontaneous vasoconstriction of blood vessels^22^. Recent studies realized highly aligned smooth muscle cell culture via precise control of the ECM^6, 8, 23, 24^. Here, we first printed rectangular stencils by DLP printer to investigate geometry-dependent vSMCs alignment. Since it is challenging to determine the alignment of the cells inside of a small fixed pattern with high cell density, we measured their nuclear orientation angle which is highly consistent with the cell body instead^25^ (**Supplementary Figure 11**). The results suggest that the percentage of aligned cells significantly increases with decreasing pattern width (**Figure 3a**). We next decreased the cell contractility by ROCK inhibitor treatment (Y27632, 10 μM, dosed 1 day after cell seeding) and the collective alignment of vSMCs in the rectangular pattern (width: 100 μm) was disrupted (**Figure 3b**). To avoid the limitation that fixed patterns restrict the free expanding of vSMCs, we printed parallel microfilaments with different filament-to-filament spacing directly on coverslips through DIW **(Supplementary Figure 12**). When the spacing between two microfilaments are less than 10 μm, cells expanded randomly across the filaments – a behavior similar to 2D culture. Increasing the spacing to 50 μm yield highly aligned vSMCs, and further increase of the spacing results in decreased degree of alignment (**Figure 3c-d**). Notably, cells that aligned inside of 50 μm spacing presented a dramatic decrease in the nuclear area and an increase of nuclear aspect ratio compared with other groups. (**Figure 3e-f**). To induce more in vivo like behavior of the vSMCs which are circumferentially aligned, we use a simple 2D model with spiral-shaped microfilaments. Indeed, we confirm cells cultured in the spiral microfilaments with 50 μm interval also presented a notable orientation guided by the printed microfilaments (**Figure 3g**).

**Figure 3.**
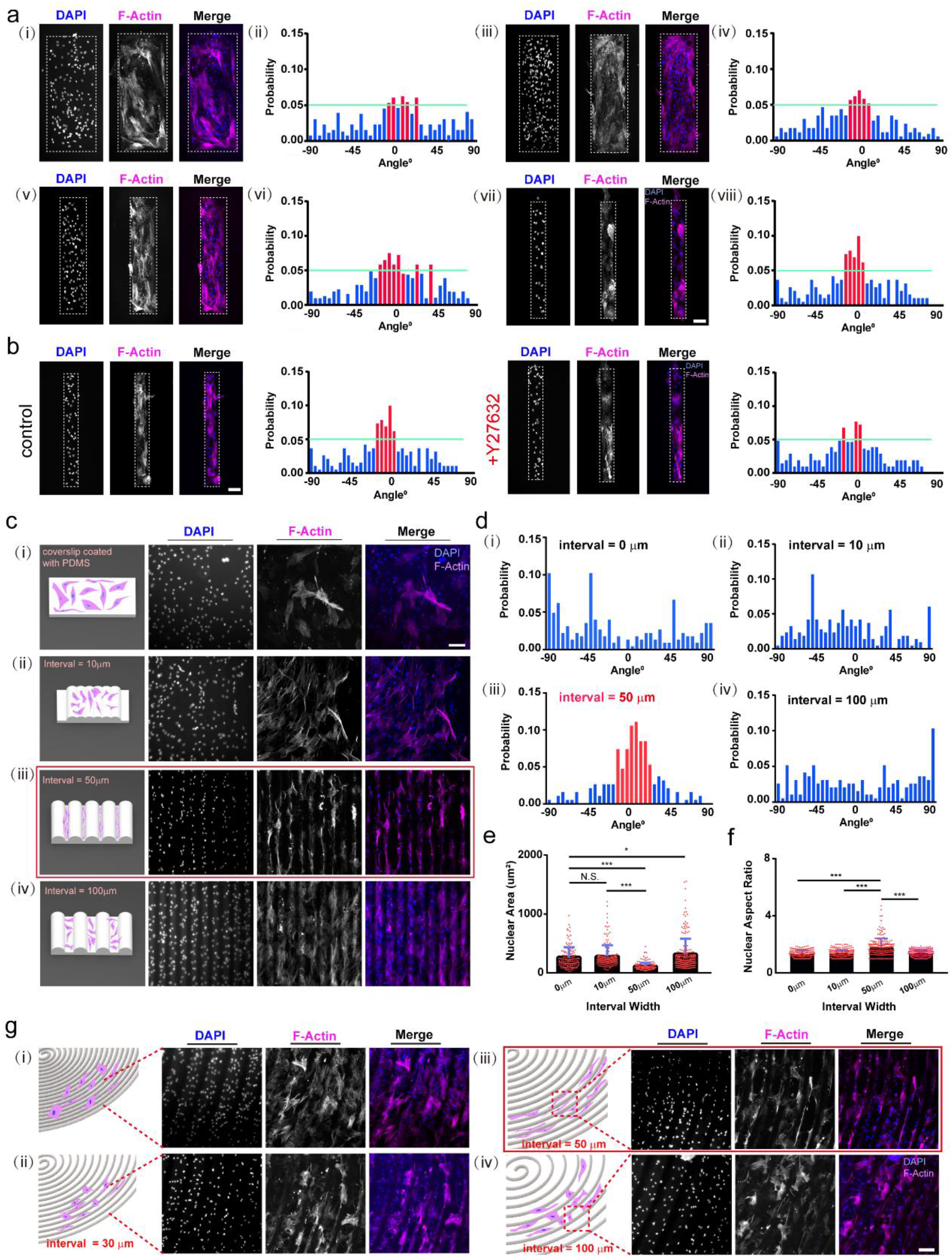
Collective alignment of vSMCs through 2D fixed rectangular patterns and direct ink writing tailored microfilaments. (a) Representative fluorescence images (i, iii, v, vii) showing vSMCs cultured on rectangular fixed patterns with decreasing widths from 400 to 100 μm and bar plots (ii, iv, vi, viii) showing the distribution of orientation angles of vSMCs in each pattern. Data plotted are mean values from at least 20 patterns for each condition. Cells were stained with phalloidin (pink) and DAPI (blue). Scale bar, 100 μm. (b) Representative fluorescence images showing vSMCs cultured on rectangular fixed patterns (width = 100 μm) with DMSO (left) and Y27632 (right) treatment and bar plots showing the distribution of oriented angles of vSMCs in each pattern. Data plotted are mean values from at least 10 patterns for each condition. Scale bar, 100 μm. (c) Sketch (left) and representative fluorescence images showing vSMCs seeded in various intervals between printed microfilaments. Cells were stained with phalloidin (pink) and DAPI (blue). Scale bar, 200 μm. (d) Bar plots showing the distribution of orientation angles of vSMCs in the increasing intervals from 0 to 100 μm. Data plotted are mean values from at least 20 patterns for each condition. Bar plots shows the differences of nuclear area (e) and nuclear aspect ratio (f) when vSMCs seeded in the increasing intervals of printed microfilaments. Data represent mean ± s.e.m. from at least three independent experiments. *, P < 0.05, **, P < 0.01, ***, P < 0.001, n.s., P > 0.05. (g) Sketch (left) and representative fluorescence images showing vSMCs seeded in the various interval between printed spiral microfilaments. Cells were stained with phalloidin (pink) and DAPI (blue). Scale bar, 200 μm.

### Realize the three-dimensional culture of vSMCs *in vitro*

Base on the planar studies that the parallel microfilaments could align vSMCs, we next demonstrate the 3D printing of circumferentially grooved vessels and 3D culturing of vSMCs for the construction of a biomimetic smooth muscle layer. While printing of the artificial vessels on a rotated rod template has been demonstrated, our current method enables a simple, one-step fabrication of complete blood vessels with a micro-grooved external surface topology for cell alignment. The resolution of the micro-grooves is at least two order of magnitude higher than previous studies which is critical for vSMC alignment. To realize both a smooth inner layer and micro-grooved external surface, we designed and 3D printed a high throughput printhead using a DLP printer ^26, 27^ **(Figure 4a-b).** The printhead is capable of simultaneously extrude 20 parallel microfilaments with fixed filament-to-filament spacing of 50 μm with high fidelity. Base on the requirements of biocompatibility, reliability, and the shear-thinning rheological properties (**Figure 4c**) for DIW, we chose PDMS 1700 (10:1 to crosslinker) as the printable ink. The printed VWMF presented a reliable mechanical property (**Figure 4d**) with a stress-strain test and exhibited mostly linear stress−strain behavior with ultimate strength up to 0.95 MPa±0.08 and failure strain up to 2.45±0.05. For vSMCs culturing on VWMF, we first coated human fibronectin (see **Method**) to facilitate the adhesion of vSMCs and then directly seed cells on the top of VWMF in a 12-well plate. Cells adhere to the VWMF after 60 minutes. We next repeated the seeding process three times every 60 minutes with a 120° rotation (**Figure 4e**). We eventually achieved a highly biomimetic smooth muscle layer with predominantly circumferentially oriented vSMCs with high viability (**Figure 4f**). (97.8%, 97.1%, and 95.65% live cells in Day 1, 2, and 3, respectively) (**Figure 4g**).

**Figure 4.**
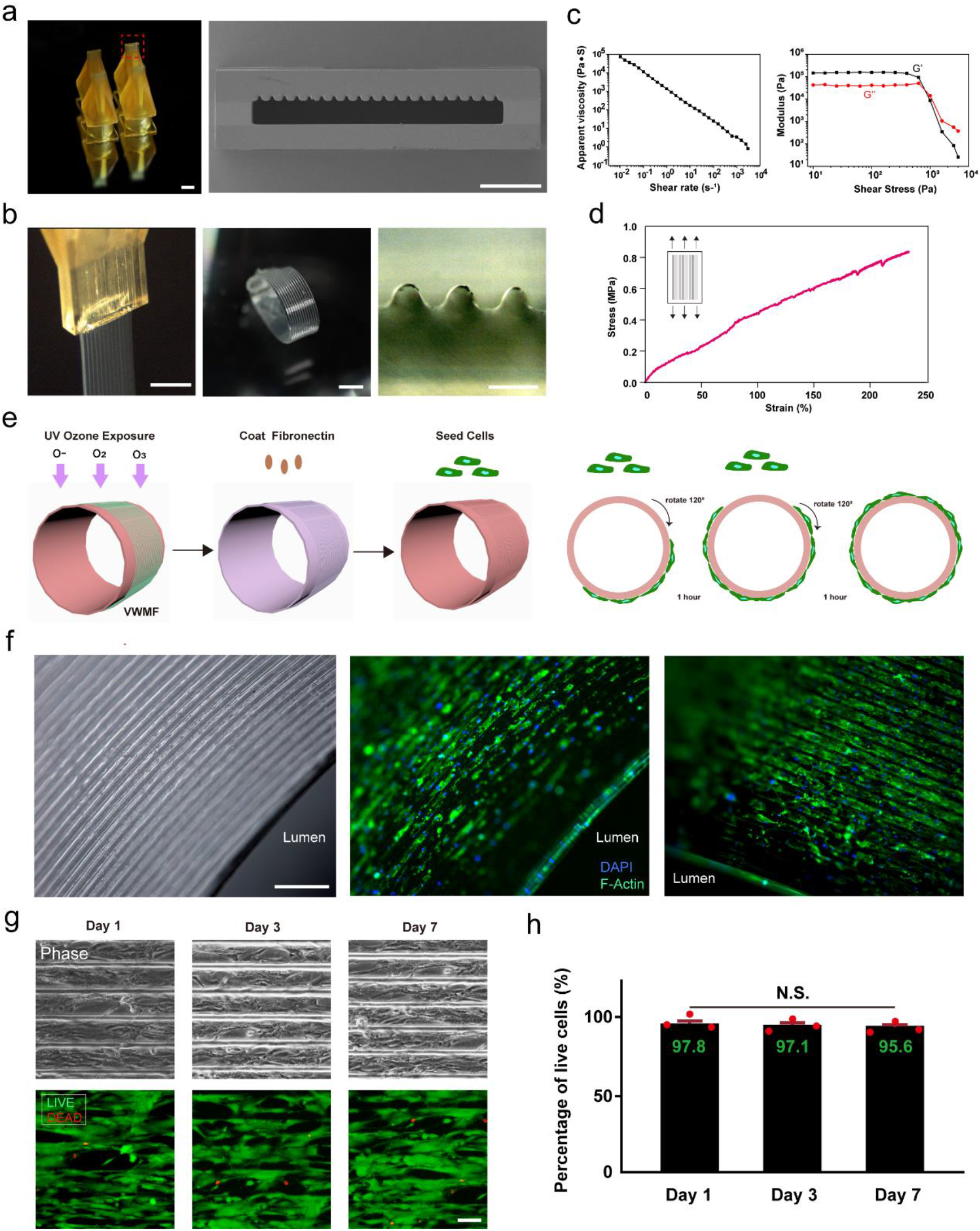
Fabricating VWMFs through a custom-printed high throughput nozzle and realizing long-term 3D culturing of vSMCs in vitro. (a) Representative images showing the custom-printed high throughput nozzles (left) and SEM images showing the cross-section of nozzles, which can extrude fixed microfilaments and vessel wall with variable thickness simultaneously (right). Scale bar, 1mm (left), 50 μm (right). (b) Representative images presenting the ink extruded from custom-printed nozzle (left), VWMF in 6-well plate (middle), and the cross-section of printed VWMF (right). Scale bar, 1 mm (left and middle), 100 μm (right). (c) Rheological properties for silicone ink. Apparent viscosity as a function of shear rate for the silicone ink (left) and shear elastic and loss moduli as a function of shear stress for the silicone ink (right). (d) Stress−strain curves of the fabricated VWMF. (e) Sketch for 3D culturing of vSMCs on VWMF. (f) Representative phase contrast image (left) and fluorescence images (right) showing the vSMCs cultured on VWMF for 5 days. Cells were stained with phalloidin (green) and DAPI (blue). Scale bar, 500 μm. (g) Representative phase and immunofluorescence images showing the LIVE / DEAD staining of vSMCs that cultured on VWMF for 1, 3, and 7 days, respectively. Scale bar, 100 μm. (h) Bar plot showing the percentage of live cells cultured on VWMF after 1, 3, 7 days, respectively. Data represent mean ± s.e.m. from at least three independent experiments. *, P < 0.05, **, P < 0.01, ***, P < 0.001, n.s., P > 0.05.

### 3D culturing inducing highly physiologically relevant phenotypes of vSMCs

In the healthy artery, vSMC shows higher contractile phenotype in contrast to synthetic phenotype, which is more often occurred in disease conditions such as atherosclerosis^28–31^. vSMC phenotype switching between these two states has long been implicated in disease progression^32^. To identify the phenotype of 3D-cultured vSMC, we examined their proliferation rate by EdU assay, which is able to detect newly generated cells. Compared to 2D-cultured vSMC on glass that had 30% newly divided cells, 3D-cultured vSMC on VWMF significantly reduced their cell proliferation rate by 18%, evidenced by EdU assay (**Figure 5a-b, Supplementary Figure 15**). These results demonstrated that curvature-induced vSMC circumneutral growth promote vSMC toward non-synthetic phenotype, which is the case *in vivo* heathy artery.

**Figure 5.**
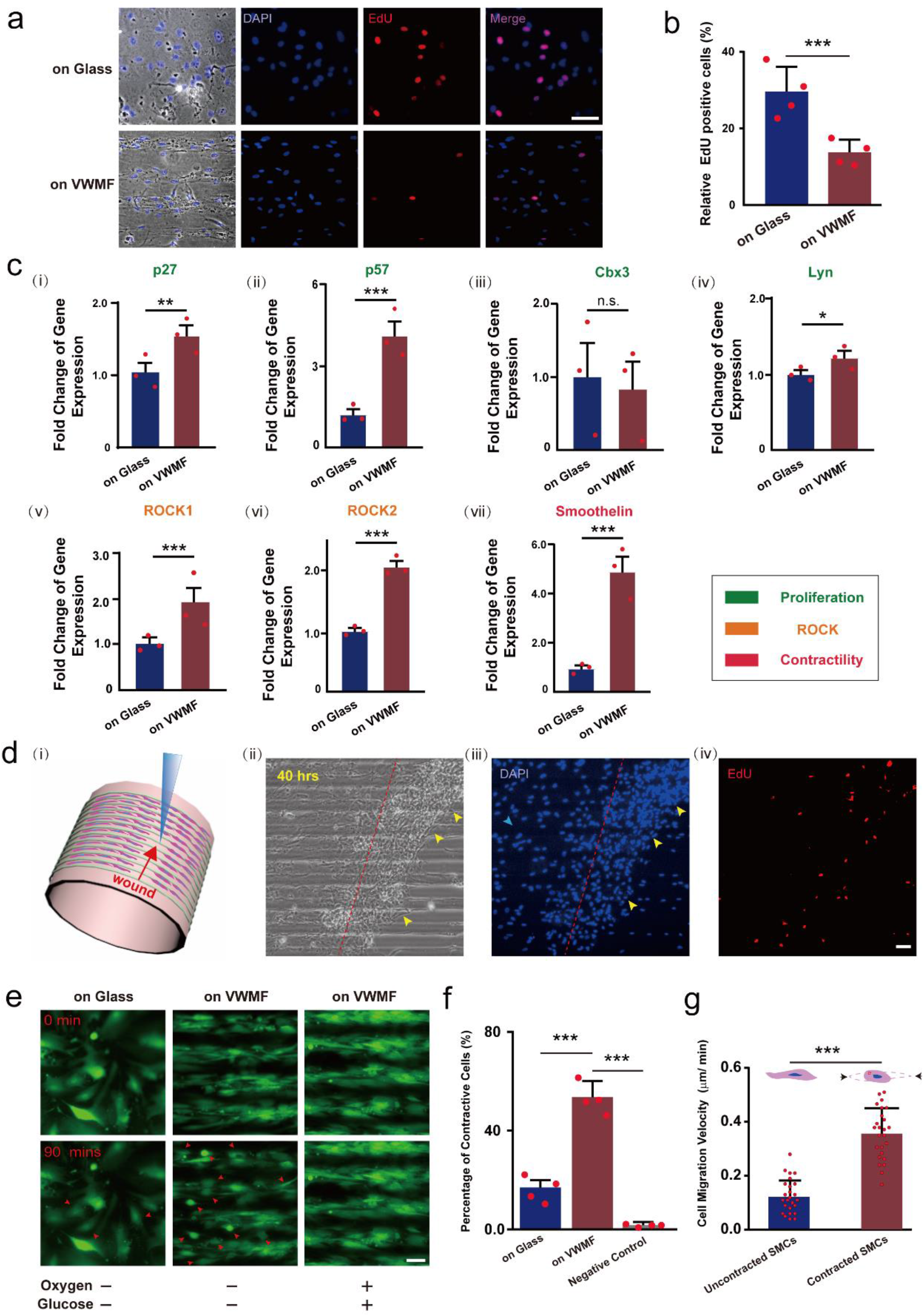
VWMF inducing a more physiological relevant phenotype and gene expression of VSMCs. (a) Representative phase contrast images (left) and fluorescence images showing the EdU test after culturing for 5 days. scale bar, 100 μm. (b) Bar plot presenting the percentage of EdU positive cells after 5 days on glass and VWMF. (c) Bar plots showing the results of phenotype markers analyzed by qPCR. Data represent mean ± s.e.m. from at least three independent experiments. *, P < 0.05, **, P < 0.01, ***, P < 0.001, n.s., P > 0.05. (d) Schematic diagram (i) indicating the wound healing assay that performed on VWMF. Representative phase image (ii) and fluorescence images (iii and iv) showing the vSMCs on VWMF, 40 hrs after generating the scratch. Cells were labeled with DAPI (blue) and EdU (red). Dotted line notes the boundary of scratch. Scale bar, 50 μm. (e) Representative fluorescence live-cell images showing the cells before and after OGD treatment. Cells were stained with CellTracker Green for better visualization. Arrowheads mark contracted cells. Scale bar, 50 μm. Bar plot showing the percentage of contracted cells during OGD (f) and the migration speed of contracted and uncontracted cells after OGD stimulation (g). Data represent mean ± s.e.m. from at least three independent experiments. *, P < 0.05, **, P < 0.01, ***, P < 0.001, n.s., P > 0.05.

To investigate the underlying molecular mechanism regarding the less proliferative phenotype, we examined the proliferation-related gene expressions. P27, P57 are reported that inhibited vSMC proliferation^33^ were significantly enhanced in 3D-cultured vSMC (**Figure 5c i-ii**). Concurrently, Cbx3 and Lyn which serve as proliferation promotional regulators were slightly downregulated and upregulated, respectively (**Figure 5c iii-iv**). ROCK signaling pathway plays a crucial role in actin cytoskeleton rearrangement and mediates various cellular manners, including cell morphology, adhesion, proliferation, migration, and contractility^34^. Through immunofluorescence staining, we observed significantly more elongated vSMCs when cultured on VWMFs, showing an average cellular aspect ratio of 5.0±2.34 vs. 1.84±0.73, respectively (**Supplementary Figure 16**). This dramatic morphological difference was further verified through the investigation of the upregulated expression of ROCK1 and ROCK2 (**Figure 5c v-vi**).To directly test whether those non-synthetic vSMC express higher contractility-related genes, we examine smoothelin^1^, which is a classical marker for mature vSMCs possessing high contractility. We indeed found that in 3D-cultured condition, smoothelin was robustly expressed (**Figure 5c vii**). These results provided strong evidence that 3D-cultured vSMC has shifted its phenotype from synthetic to contractile status, which suggests our system could provide more *in vivo*-like microenvironment for vSMCs.

An important function of vSMCs is to regulate blood flow mainly through vasodilation and vasoconstriction mediating^35^. However, the normal phenotypes of vSMCs can be modulated by environmental cues and lead to related diseases. For example, the aberrant proliferation of vSMCs after vascular damage was identified as the initiation and progression in most cases of early atherosclerosis^36^. To mimic this pathological model, we created scratches on VWMF after 7-day culturing (**Figure 5d i**). We note that the cells close to the wound proliferated faster than the cells distant from the wound (**Figure 5d ii-iv**) whereas the planarly cultured cells proliferate only moderately (**Supplementary Figure 17**). Lastly, we aim to investigate how 3D-cultured vSMC responds under pathological conditions comparing to 2D-culture vSMC. We created an ischemia-like environment by oxygen and glucose deprivation (OGD) from the culturing systems. Using time lapse imaging, surprisingly, we found vSMCs on VWMF show remarkably higher contractility than 2D-cultured ones, both subject to OGD. Specifically, in the first 90 minutes, more than 53.5% of the 3D cultured vSMCs on VWMF presented a significant contraction, showing a notable reduction of cellular aspect ratio, compared to only 16.9% of 2D cultured vSMCs on glass showing contraction (no notable changes for the negative group with 1.7%) (**Figure 5e-f**). We next performed a long-term imaging of the contacted cells to eliminate the possibility that those round-up vSMCs were enduring the process of apoptosis. In the following 10 hrs, nearly all the contracted cells kept alive (**Supplementary Figure 18**), and some of them even re-expended on the substrate (**Supplementary Figure 19**). Interestingly, we observed that those highly contractile vSMCs were more dynamic with the migration speed of 0.356±0.09 μm/min than the diastolic vSMCs with 0.125±0.06 μm/min (**Figure 5g**).

Altogether, our results demonstrated that 3D-culture vSMCs exhibited remarkable differences in cell divisions, gene expressions and their contractility upon pathological stimulation. Our demonstrated 3D culturing method allows vSMCs grow and function in *in vivo*-like environment, which would be of great significance for further biological study *in vitro*.

## DISCUSSION

High-resolution 3D printing enables the *in vitro* reconstruction of clinically relevant vSMCs microenvironment, which could potentially lead to significant advancement in biomedical research field. Harnessing multiple 3D printing technologies, we first manufactured various types of ECMs with complex geometries and induced more behaviors of vSMCs than from conventional 2D cell culture on petri dishes. Besides, narrowed space facilitating the spontaneous alignment of vSMCs, was validated by direct printing closed parallel microfilaments with different interval from 10 μm to 100 μm. Further employing customized printhead, we demonstrate a novel 3D printing method to rapidly manufacture in vitro vessel models for 3D culture of vSMCs. We found that the precisely engineered circumferentially grooved surface topology of the vessels direct circumferential orientation of vSMCs. The *in vivo*-like phenotype was further verified by testing the expression of typical markers. Lastly, we found that 3D cultured vSMCs on VWMF exhibit a collectively higher degree of contraction than 2D cultured SMCs when incubated in a simulated pathological ischemic environment, and the contracted cells are able to transform into motile phenotypes. Despite the successfully aligned vSMCs, we do not observe the self-contraction of VWMF that induced by vSMCs’ spontaneous contraction due to high elastic modulus of the printed vessel materials. Development of 3D in vitro culture vSMCs using new 3D printable, biocompatible and degradable materials^37^ are underway to investigate artificial vessel models capable of spontaneous contraction. Through 3D printing technologies and the investigations on physiological relevant phenotypes of SMCs induced *in vitro* demonstrated in this work, we provide a versatile in vitro platform to facilitates in vitro vascular reconstruction and drug screen research.

## ACKNOWLEDGEMENT

The authors thank Dr. Zhu Zhu for providing the sample of artery vessel of NG2DsRed. The authors are grateful for the financial support of this research by the National Natural Science Foundation of China (No. 51905446). Authors would like to acknowledge funding support from the Westlake University and Bright Dream Joint Institute for Intelligent Robotics. We also thank the Microscopy Core Facility of Westlake University for providing advice and assistance.

## AUTHOR CONTRIBUTIONS

P.Z., X.L.,Y.S., J.J., and N.Z. designed experiments; P.Z., X.L., W.X, M.W., and H.Z. performed experiments; P.Z. and H.C. designed the print nozzle and set up the platform; P.Z., M.W., W.X., J.L., and C.Y. analyzed data; P.Z., X.L, W.X., J.J, and N.Z. wrote the manuscript. All authors edited and approved the final manuscript.

## COMPETING INTERESTS

The authors declare no competing financial interest.

## Notes

### Competing Interest Statement

The authors have declared no competing interest.

