## Supplementary material for "Inducing highly physiologically relevant phenotypes of human vascular smooth muscle cells via 3D printing": Supplemetary Information

**SUPPORTING INFORMATION**

### Inducing highly physiologically relevant phenotypes of human aortic vascular smooth muscle cells in vitro via 3D bioprinting

*Peiran Zhu[1,2,3#], Xuzhao Li[4,5#], Wang Xin[1,2,], Menglin Wang[1,2], Chengzhen Yin[1,2], Jinze Li[4,5], Hangyu Chen[1,2], Hengjia Zhu[1,2], Yubing Sun[6,7], Jiemin Jia[4,5*] and Nanjia Zhou[1,2*]*

#### ADDRESS

[1]Key Laboratory of 3D Micro/Nano Fabrication and Characterization of Zhejiang Province, School of Engineering, Westlake University, 18 Shilongshan Road, Hangzhou 310024, Zhejiang Province, China.

[2]Institute of Advanced Technology, Westlake Institute for Advanced Study, 18 Shilongshan Road, Hangzhou 310024, Zhejiang Province, China.

[3] School of Materials Science and Engineering, Zhejiang University, Hangzhou, China.

[4]Key Laboratory of Growth Regulation and Translation Research of Zhejiang Province, School of Life Sciences, Westlake University, Hangzhou, Zhejiang Province, 310024, China.

[5]Institute of Biology, Westlake Institute for Advanced Study, Hangzhou, Zhejiang Province, 310024, China.

[6]Department of Mechanical and Industrial Engineering, University of Massachusetts, Amherst, Massachusetts 01003, USA.

[7]Department of Chemical Engineering, University of Massachusetts, Amherst, Massachusetts 01003, USA.

#These authors contribute equally to this work.

#### Supplementary Information:

##### Supplementary Figures and Captions

##### Supplementary Table

##### Methods

##### References

#### Supplementary Figures and Captions

#### Supplementary Figure 1

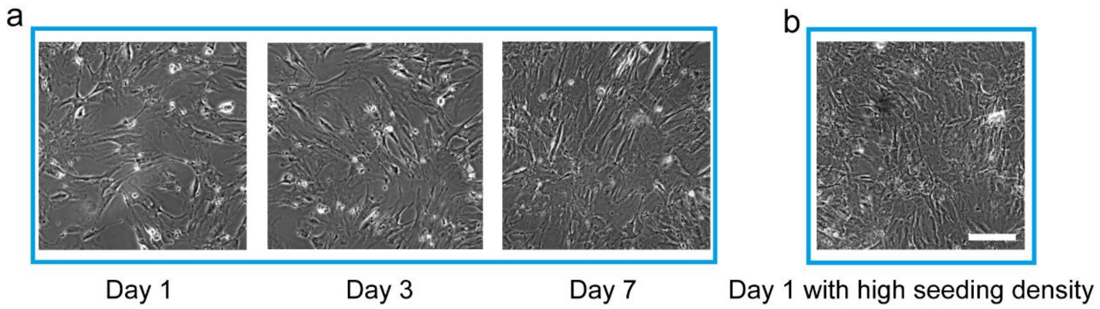


**Supplementary Figure 1.** (a) Representative phase contrast images showing VSMCs cultured on glass for 1, 3, and 7 days. Although cells came to more than 90% conflunence, no spontaneous orientation of VSMCs can be observed. And we also tried to seed a high cell concentration (0.4 million cells/ml) from the beginning and no orientation after 3 days as well (b). Scale bar, 100 μm.

#### Supplementary Figure 2

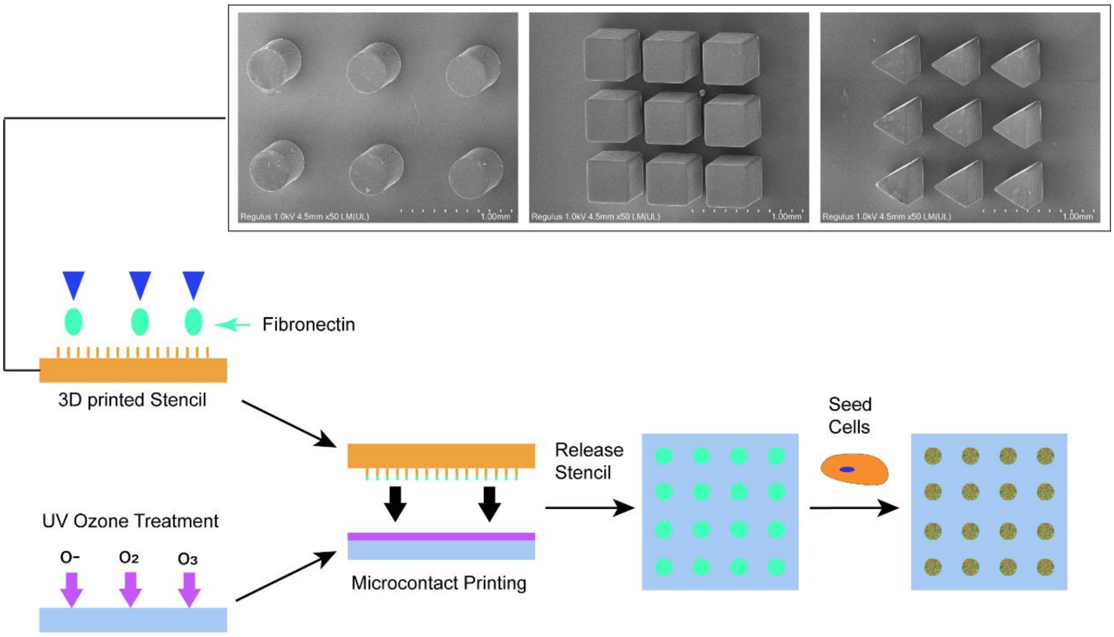


**Supplementary Figure 2.** Schematic diagrams showing the process of microcontact printing. 3D printed stamps were soaked in fibronectin solution (50 μg·ml$^{-1}$ in sterile, deionized water) for 1 h. A fibronectin coated stamp was gently placed on the top of the flat PDMS substrate, after treating with UV ozone for 7 min. The stamp was pressed gently to facilitate the transfer of fibronectin to coverslips. Coverslips were rinsed with PBS and transferred to standard 12-well tissue culture plates for seeding cells.

#### Supplementary Figure 3

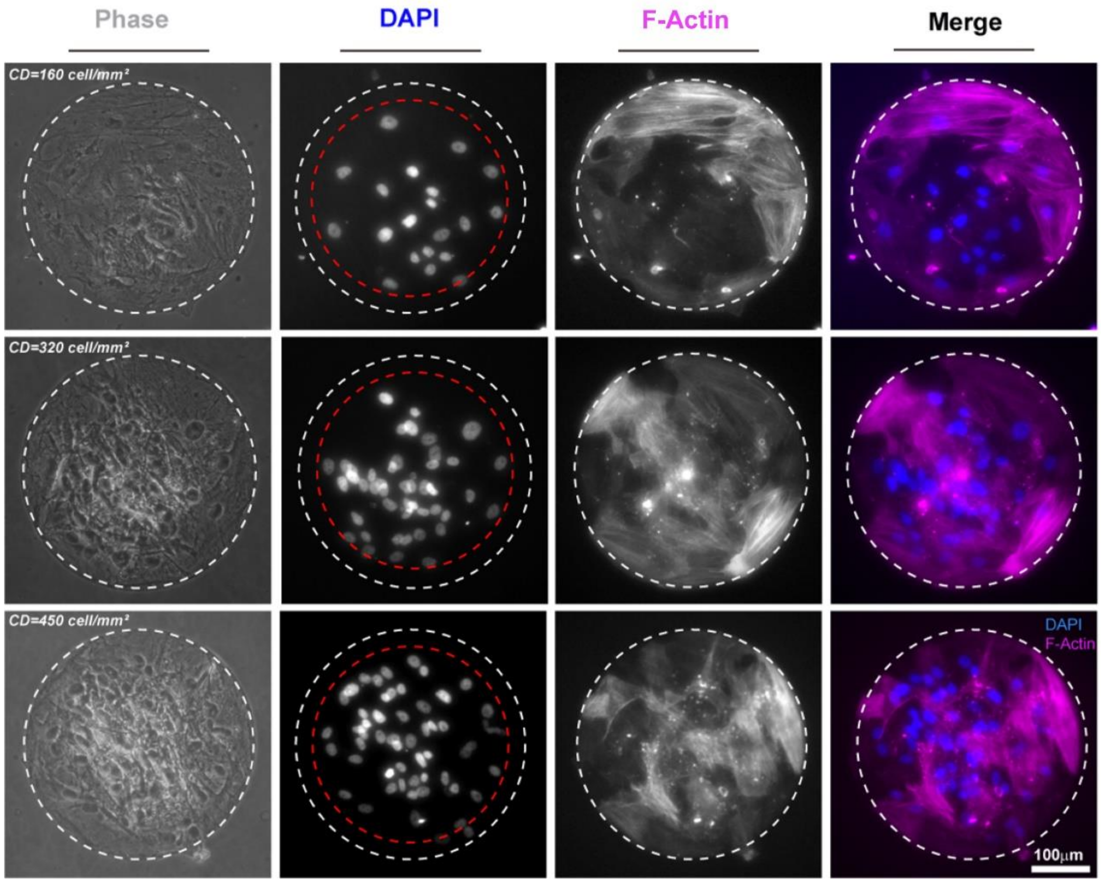


**Supplementary Figure 3.** Representative phase contrast (left row) and fluorescence images (violet shows phalloidin staining, blue shows nuclei staining). vSMCs spontaneously crowded in the central of the circular fixed pattern 2 day after seeding. And the start seeding density would not affect the behavior. Scale bar, 100 μm.

#### Supplementary Figure 4

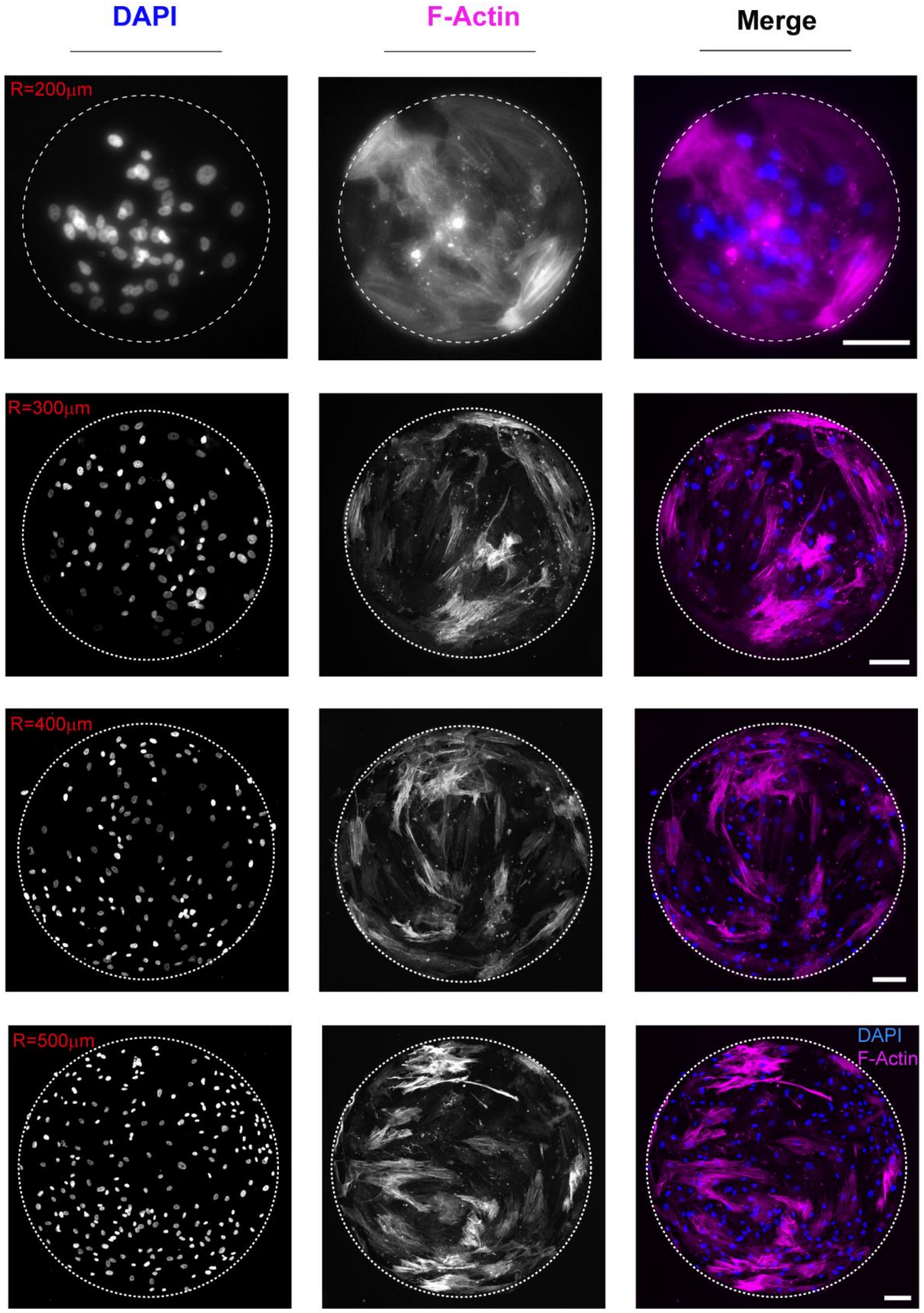


**Supplementary Figure 4.** Fluorescence images showing vSMCs cultured on circular fixed patterns with an increasing diameter, 400 μm, 600 μm, 800 μm, 1000 μm, respectively. Cells were stained with phalloidin (violet) and DAPI (blue). vSMCs distributed more uniformly with the increasing diameter of fixed patterns. Scale bar, 100 μm.

#### Supplementary Figure 5

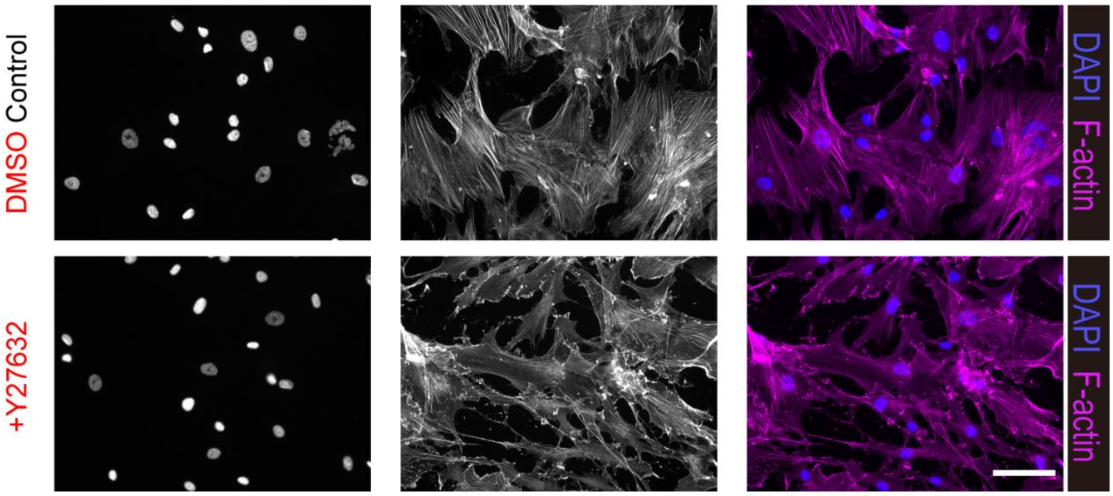


**Supplementary Figure 5.** Fluorescence images showing vSMCs treated with 10 μM DMSO (control) and 10 μM Y27632 on glass. Cells were stained with phalloidin (violet) and DAPI (blue). Scale bar, 100 μm.

#### Supplementary Figure 6

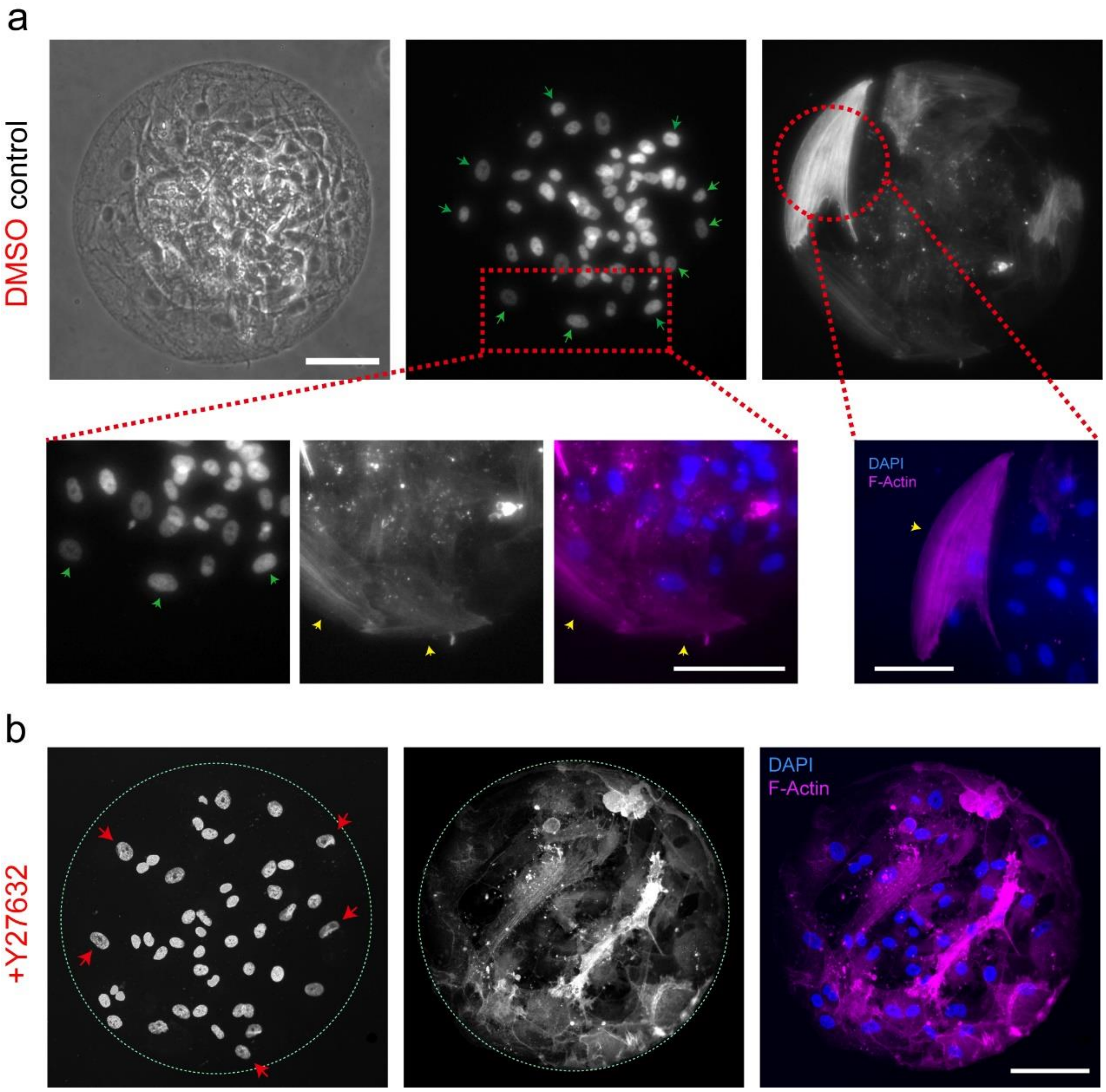


**Supplementary Figure 6.** (a) Fluorescence images showing vSMCs crowded in the central of the fixed patterns. While cells distributed at the periphery are highly expended, and both nuclear and cytoskeleton perfectly fitted the curvature of periphery, previous manners of cells at the periphery were disrupted after ROCK inhibitor treatment (b). Cells were stained with phalloidin (violet) and DAPI (blue). Scale bar, 100 μm

#### Supplementary Figure 7

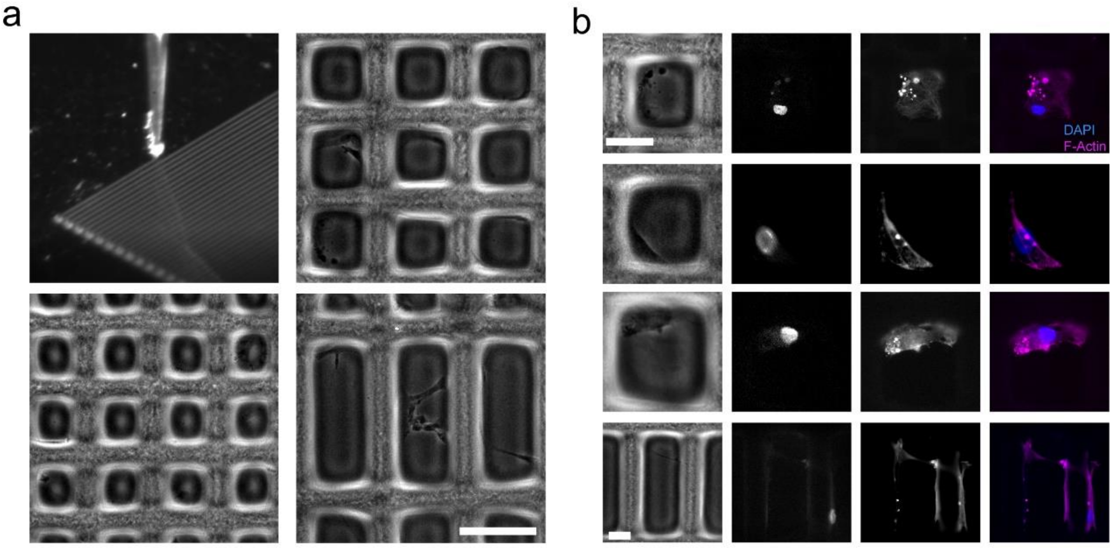


**Supplementary Figure 7.** (a) Representative phase contrast images showing the processing of 2D scaffold printing. (b) Representative fluorescence images show the vSMCs cultured on 2D scaffold. Cells were stained with phalloidin (violet) and DAPI (blue).Scale bar, 50 µm.

#### Supplementary Figure 8

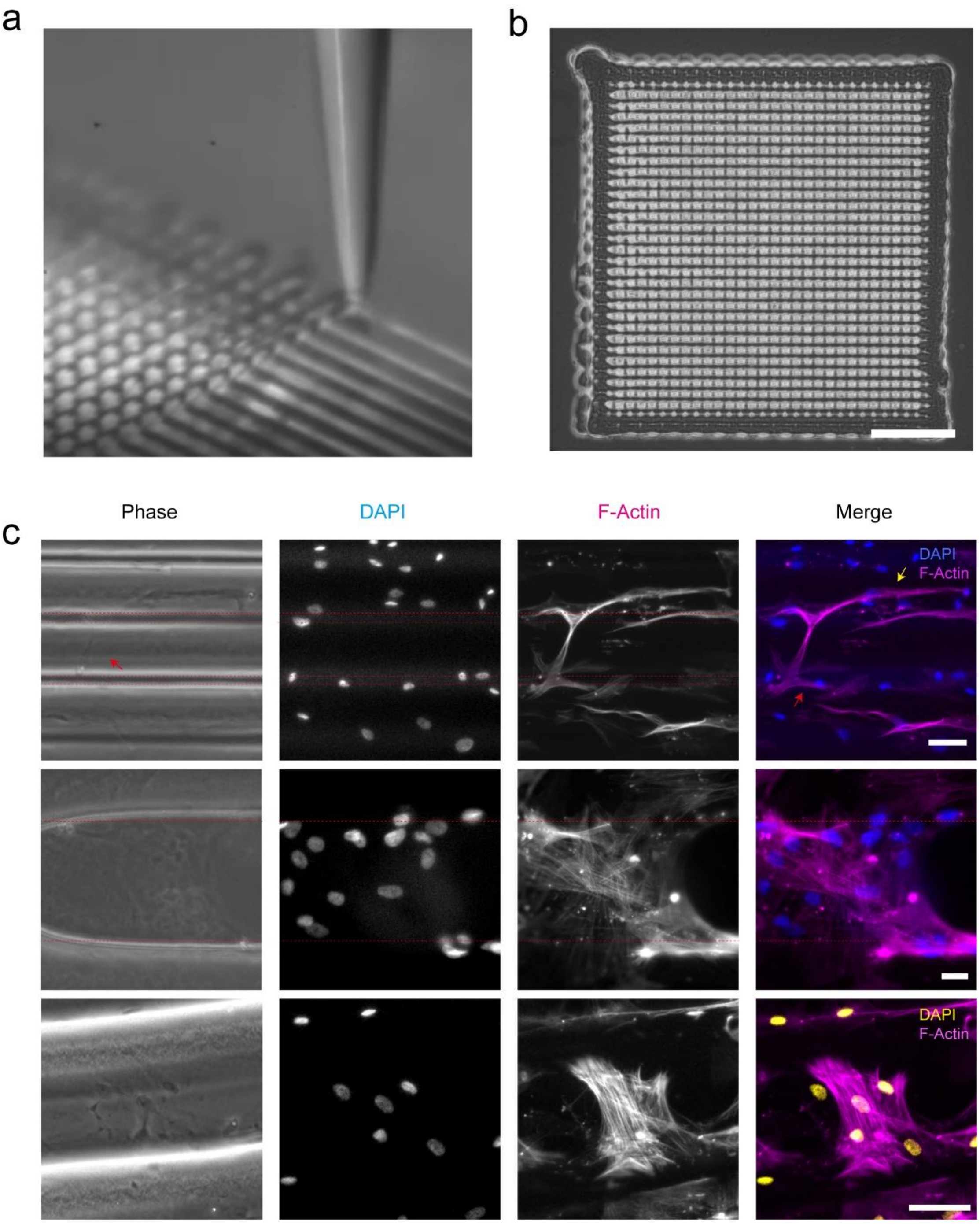


**Supplementary Figure 8.** (a)-(b) Representative phase contrast images showing the processing of 3D scaffold printing. Scale bar, 500 µm (c) Representative fluorescence images showing the vSMCs cultured on 3D scaffold with diverse behaviors. Cells were stained with phalloidin (violet) and DAPI (blue and yellow). Scale bar, 50 µm.

#### Supplementary Figure 9

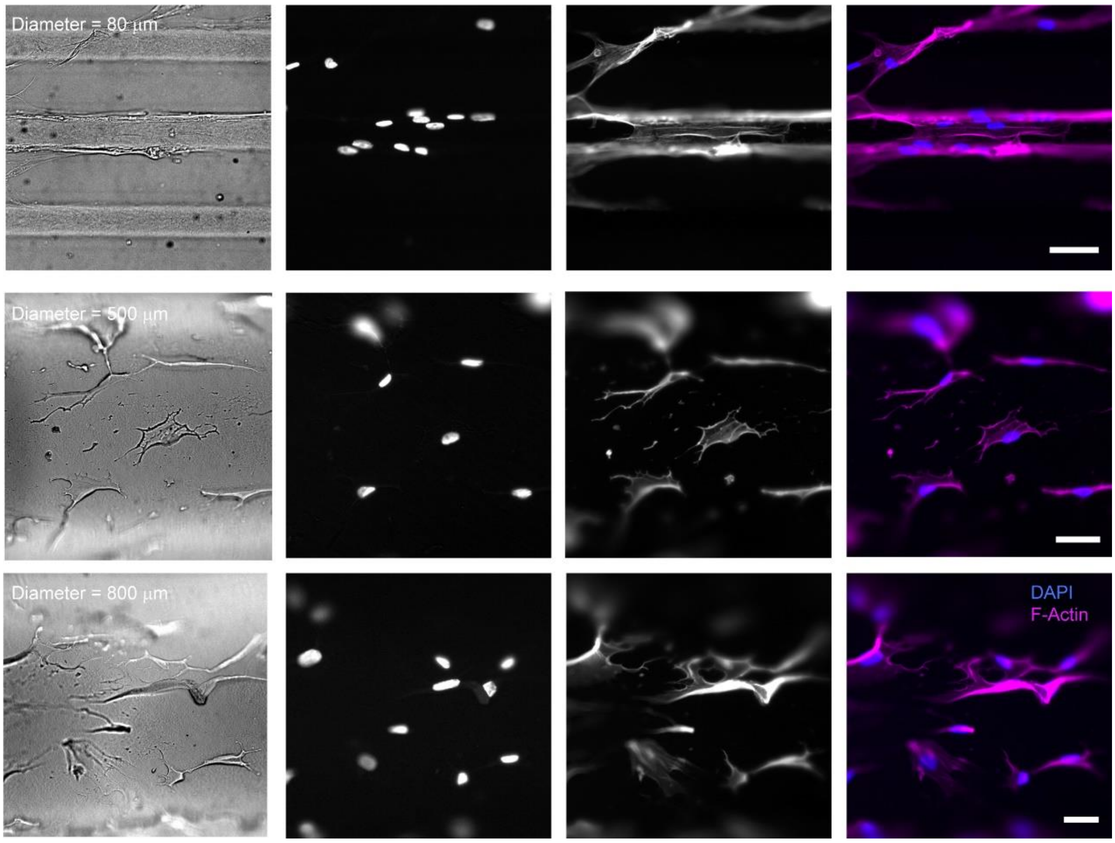


**Supplementary Figure 9.** Representative phase contrast and fluorescence images showing vSMCs cultured on the top surface of the microfilaments with increasing diameter. Cells were stained with phalloidin (violet) and DAPI (blue). Scale bar, 100 μm.

#### Supplementary Figure 10

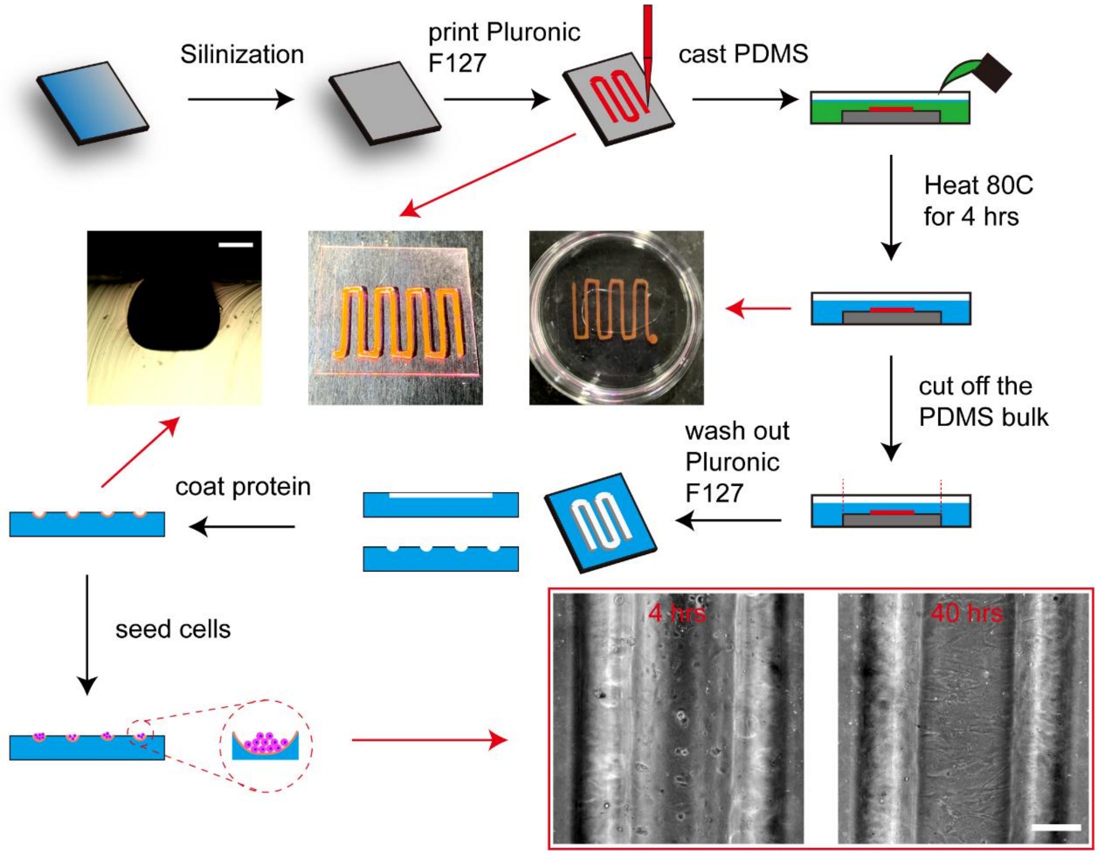


**Supplementary Figure 10.** Schematic diagram showing the fabrication of concave ECM for vSMCs culturing. In short, we first printed fugitive materials on the coverslip to make a negative mold and then poured PDMS. After incubating 2 hours under 80 $^{o}$C for PDMS curing, the fugitive material would be washed away from the PDMS and left concave channels. vSMCs would be spontaneously oriented along circumferential direction after 40 hours. Scale bar, 800 µm.

#### Supplementary Figure 11

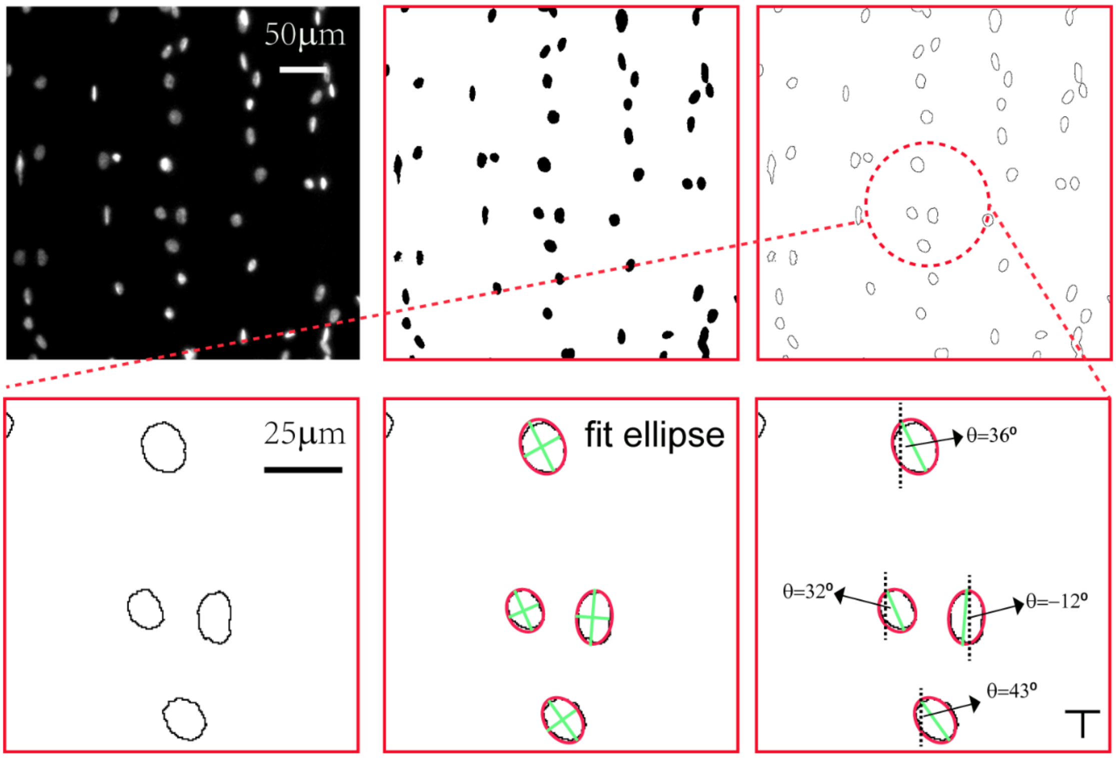


**Supplementary Figure 11.** Schematic diagram showing the strategy for orientation angle measurement.

#### Supplementary Figure 12

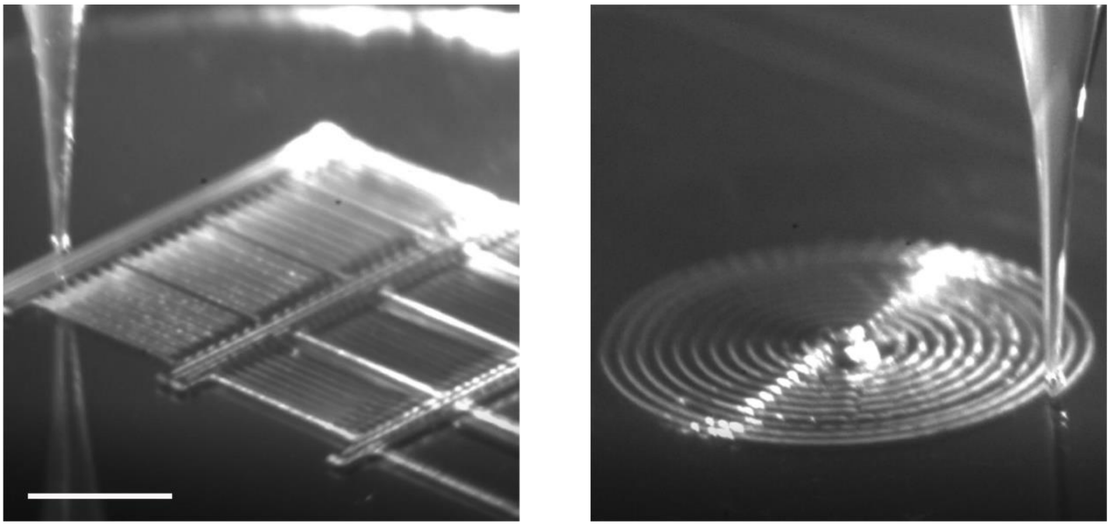

**Supplementary Figure 12.** Representative phase images showing the printing processing of parallel (left) and spiral (right) microfilaments. Scale bar, 1 mm.

#### Supplementary Figure 13

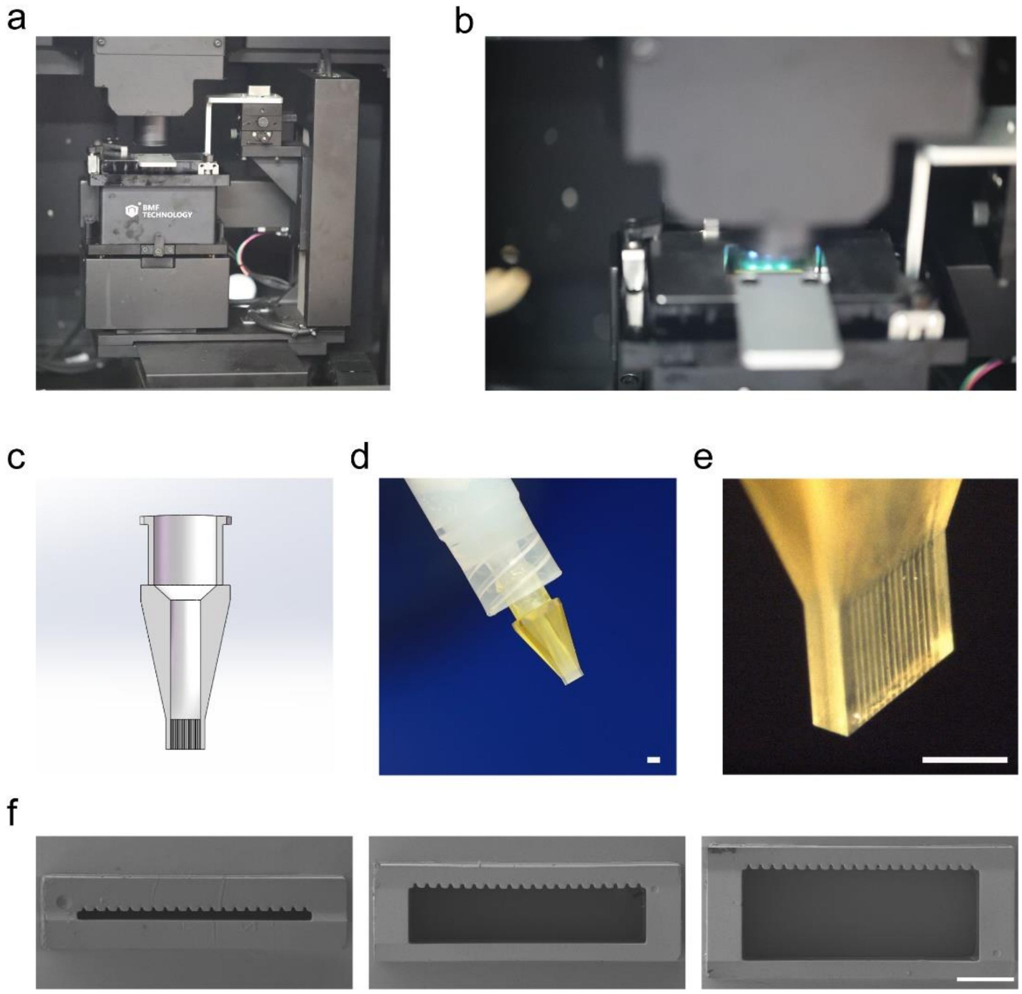


**Supplementary Figure 13.** Representative images showing the fabricate process of customized nozzle through a DLP printer (a-b). The design of high throughput nozzle (c) and then mounted on a syringe (d-e). (f) SEM images showing the cross section of high throughput nozzles. Scale bar, 500 μm.

#### Supplementary Figure 14

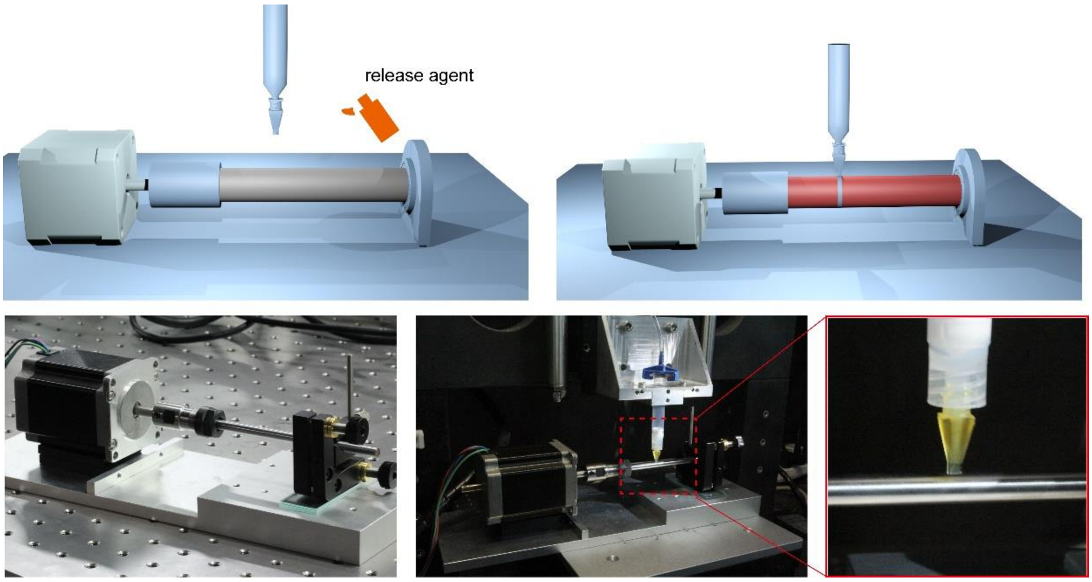


**Supplementary Figure 14.** Schematic diagrams showing the fabrication process of VWMF and the setup construction.

#### Supplementary Figure 15

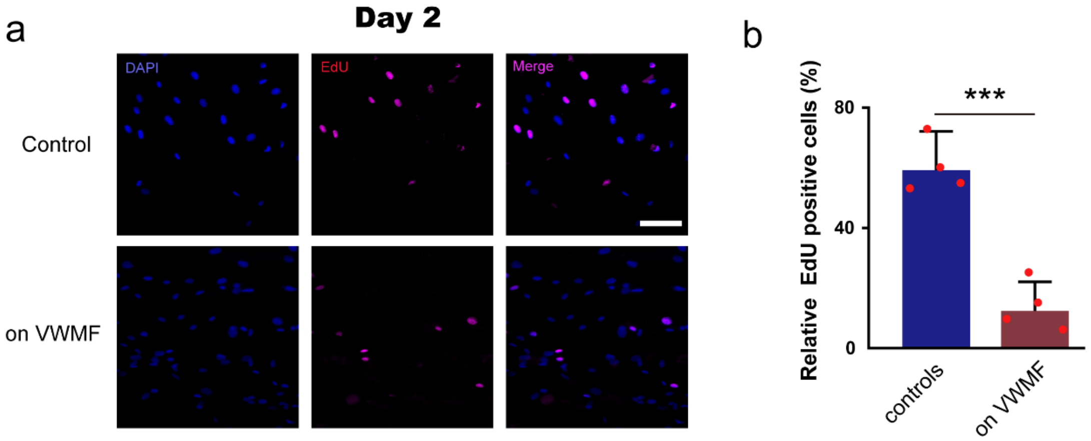


**Supplementary Figure 15.** (a) Representative fluorescence images showing the EdU test after culturing for 2 days. scale bar, 100 μm. (b) Bar plot presenting the percentage of EdU positive cells after 2 days on glass and VWMF. Data represent mean ± s.e.m. from at least three independent experiments. *, P < 0.05, **, P < 0.01, ***, P < 0.001, n.s., P > 0.05.

#### Supplementary Figure 16

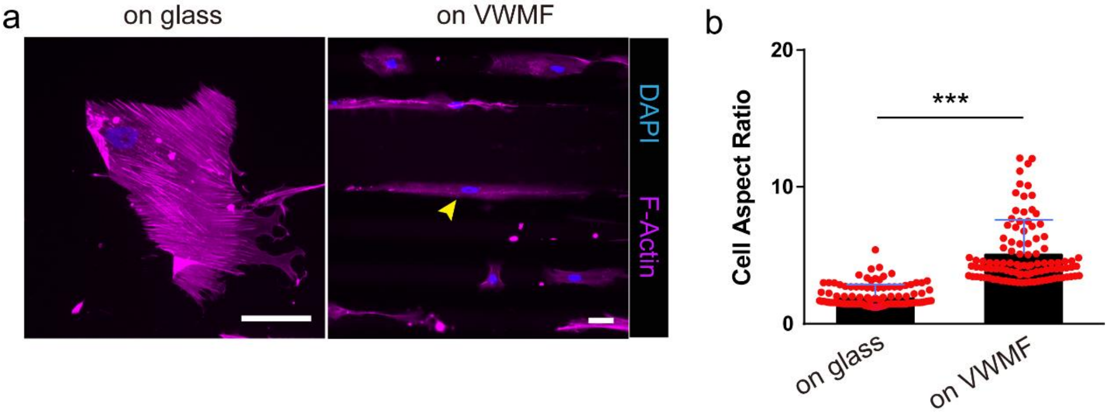


**Supplementary Figure 16.** (a) Representative fluorescence image showing the vSMCs cultured on glass (left) and VWMF (right). scale bar, 50 μm. Cells were stained with phalloidin (violet) and DAPI (blue). (b) Bar plot presenting the cellular aspect ratio on glass and VWMF. 50 cells for each condition. Data represent mean ± s.e.m. from at least three independent experiments. *, $P < 0.05$, **, $P < 0.01$, ***, $P < 0.001$, n.s., $P > 0.05$.

#### Supplementary Figure 17

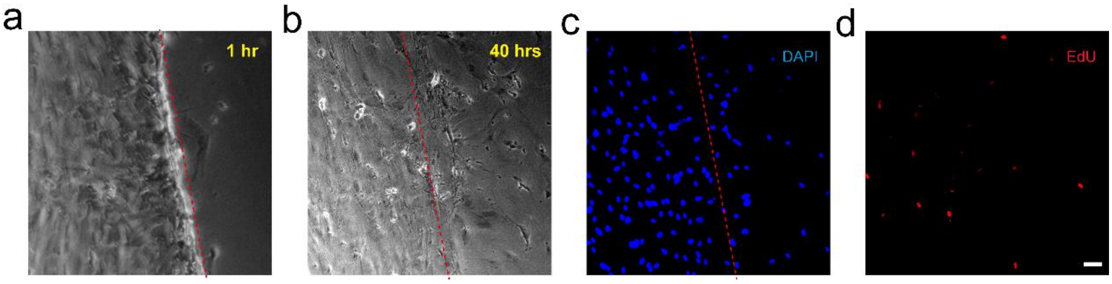


**Supplementary Figure 17.** Representative phase images and fluorescence images showing the cells planar cultured on glass (a) and 40 hours (b) after generating the scratch. Cells were labeled with DAPI (c) and EdU (d). Dotted line notes the boundary of scratch. Scale bar, 50 µm.

#### Supplementary Figure 18

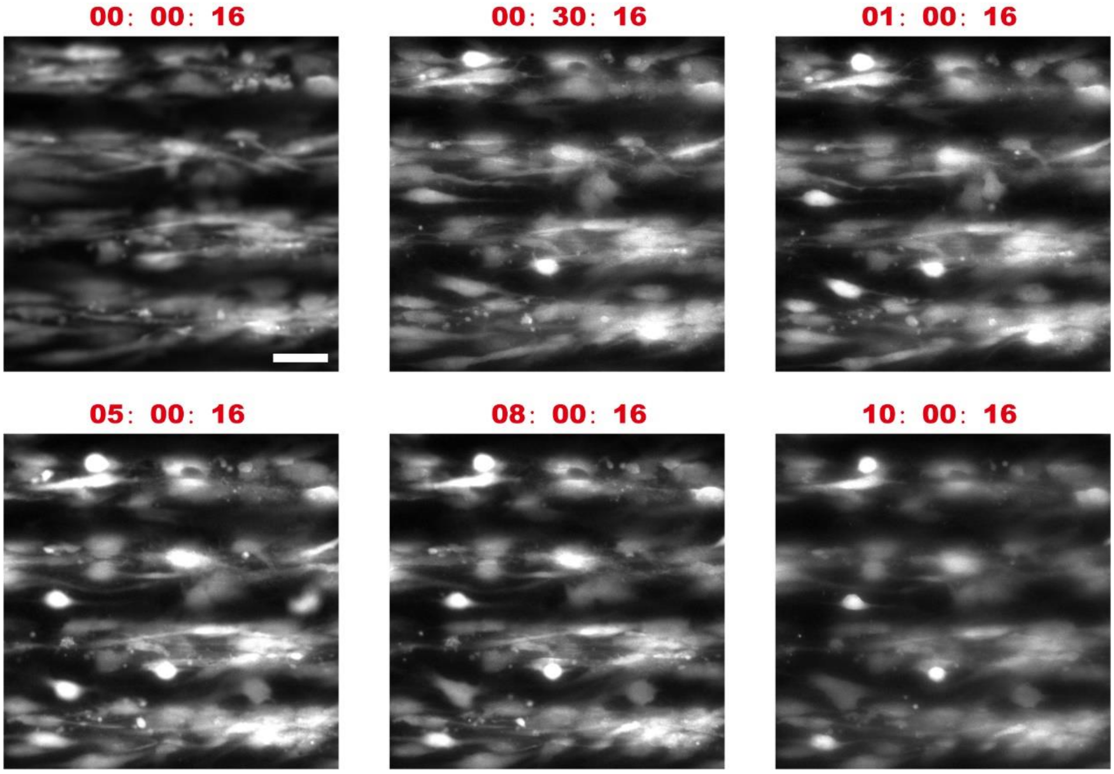


**Supplementary Figure 18.** Representative Grayscale images showing the time-lapse images of vSMCs that cultured on VWFM after OGD stimulation. Cells were stained with CellTracker Green for better visualization. Although sharply contracted at first hour, the vSMCs could keep great viability for the following ten hours in the OGD environment. Scale bar, 50 µm. Time scale, Hours: Minutes: Seconds.

#### Supplementary Figure 19

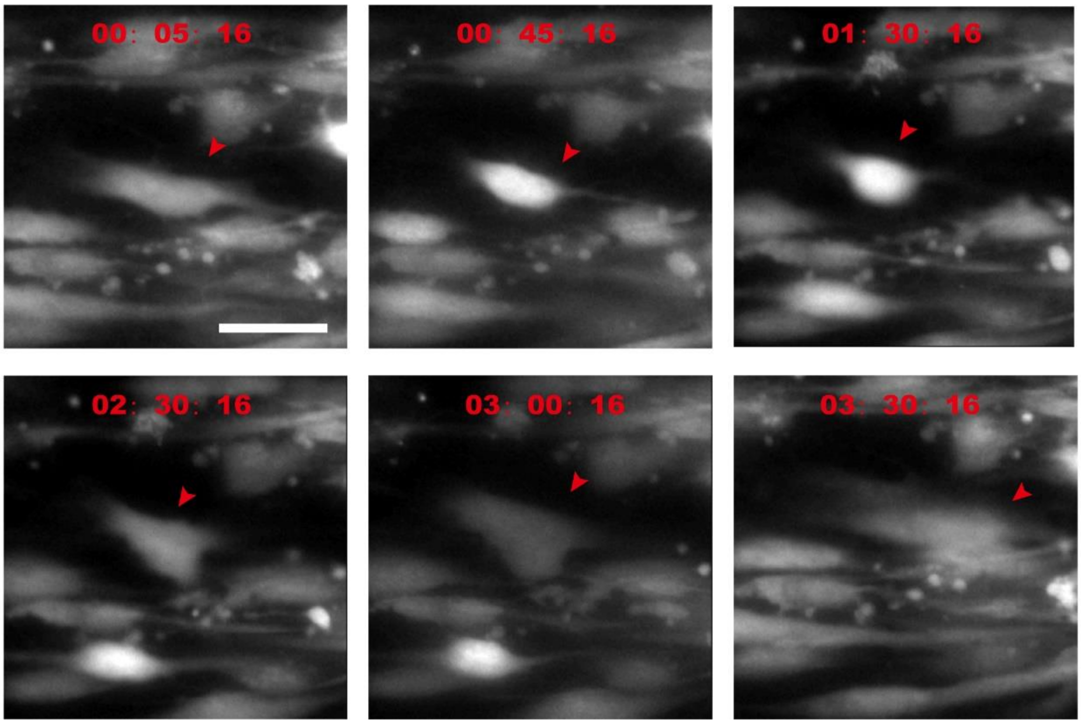


**Supplementary Figure 19.** Representative Grayscale images showing the time-lapse images of vSMCs that cultured on VWFM after OGD stimulation. Cells were stained with CellTracker Green for better visualization. The highly contracted vSMC could re-adhere to the matrix within three hours. Scale bar, 50 μm. Time scale, Hours: Minutes: Seconds.

#### Supplementary Table 1: Primers for qRT-PCR

| Gene | Forward | Reverse |
|---|---|---|
| GAPDH | GGAGTCCACTGGCGTCTTCA | GTCATGAGTCCTTCCACGATACC |
| P27 | TAATTGGGGCTCCGGCTAAC | GAAGAATCGTCGGTTGCAGGT |
| P57 | GGGCCTCTGATCTCCGATTT | GTTCCCTCGGGGCTCTTTG |
| Cbx3 | CCACGCCGCGAACGTAATA | TTCAGGCTCTGCCTCTTCAA |
| Smoothelin | AAAGAGCTAAGCCGAGCCTG | CGTCGGTTCCTTTCTGGTGA |
| ROCK1 | AAGAGGGCATTGTCACAGCA | AGCATCCAATCCATCCAGCA |
| ROCK2 | AACGTCAGGATGCAGATGGG | CAGCCAAAGAGTCCCGTTCA |
| Fyn | GGAAAAAGACCAGTCCTGCCT | TCCCTCTCCTCCGTCAGTTT |
| Lyn | TAAACAGCAAAGGCCAGTTCC | GGGTGGATGCCATCATAGGG |
| ADAMTS7 | TGTGTCAACACCCAGACAGG | CTCACAGCGGGGAGGCT |
| MYH8 | ATAGCAGCGCGAAGAAAGGT | TTGCCCCAGGAGTTTTGGTT |

#### METHOD

##### 1. Cell culture

The Human Aortic Smooth Muscle cells (T/G HA-VSMC) were grown in F-12K medium supplemented by 1% penicillin/streptomycin (Gibco); 0.05 mg/ml ascorbic acid; 0.01 mg/ml insulin; 0.01 mg/ml transferrin; 10 ng/ml sodium selenite; 0.03 mg/ml Endothelial Cell Growth Supplement (ECGS); fetal bovine serum to a final concentration of 10%, HEPES to a final concentration of 10 mM, TES to a final concentration of 10 mM. Cells were cultured at 37°C and with 5% $CO_2$. Media were changed every three days, and cells were passaged when they were nearly 90% confluent using 0.25% trypsin-EDTA (Gibco).

##### 2. Microcontact printing

Stencils were generated using a high-resolution DLP 3D printer to create stamps and microcontact printing techniques, as described previously[1-3]. Briefly, stamps were created using SolidWorks and sliced. The sliced layers of exposure images were then loaded into a DLP 3D printer (nanoArch P140, BMF Material Technology Inc., Shenzhen, China). Round glass coverslips with a diameter of 18 mm (Fisher Scientific) were spin-coated (Spin Coater; Laurell Technologies) with a thin layer of PDMS prepolymer containing PDMS base monomers and curing agents (10:1 w/w; Sylgard 184, Dow-Corning) before thermal curing at 110°C for at least 24 hrs. 3D printed stencils were soaked in fibronectin solution (50 $\mu g \cdot ml^{-1}$ in sterile, deionized water) for 1 hr. Excess fibronectin was then washed away by DI water, and the stamps were dried with a stream of $N_2$. Fibronectin coated stamps were gently placed on top of the flat PDMS substrates, after treating with UV ozone for 7 min. The stamps were pressed gently to facilitate the transfer of fibronectin to coverslips. Protein adsorption to all PDMS surfaces not coated with fibronectin was prevented by immersing coverslips in 0.2% Pluronics F127 NF solution (Sigma) for 30 min. Coverslips were rinsed with PBS and transferred to standard 12-well tissue culture plates for seeding cells.

##### 3. 2D/3D scaffolds fabrication

Direct ink writing was used to manufacture 2D and 3D scaffolds. First, the printable ink was prepared by combining a 10:1 weight ratio of SE1700 base: curing agent (Dow

Corning). Components were mixed in a planetary centrifugal mixer (Thinky ARE-310) at 2000 rpm for 6 min, and subsequently loaded into a 3cc syringe barrel (EFD Inc., East Providence, RI, USA). The syringe with degassed ink was mounted on a custom-built 3D printing setup; the ink was extruded through steel or glass microcapillary nozzles with 25-500 μm inner diameter. A pneumatic dispenser (Ultimus V, Nordson EFD) was used to extrude the ink. The 2D/3D scaffolds were printed on a 25 mm coverslip (Thermo Fisher) and then incubated under 80 °C for 2 hrs. The printed scaffolds were sterilized by UV exposure and ethanol immersion before cell culture.

**4. Cell viability assays**

Cells were washed with PBS twice before staining and then treated with LIVE/DEAD reagent, including calcein (1 μl/ml), ethidium homodimer-1 (EthD-1, 4 μl/ml), and Hoechst solution (0.5 μl/ml). After incubated at 37°C for 30 min, LIVE/DEAD images were acquired on a fluorescence microscope.

**5. Immunofluorescence staining**

Samples were fixed with 4% paraformaldehyde (PFA) and permeated with 0.1% Triton-X. Then the samples were blocked with 3% bovine serum albumin (BSA) in phosphate buffered saline (PBS), and subsequently incubated with primary antibodies and secondary antibodies in the blocking buffer at room temperature.

**6. Rock Inhibitor Treatment**

For Y27632 (MCE, Shanghai, China) treatment, a stock solution (10 mM in DMSO) was prepared and diluted in culture media to 10 μM. Cells were treated 1 h after cell seeding and the same volume of DMSO was added into control group.

**7. High throughput nozzle fabrication**

The high throughput nozzles were designed using a 3D computer-aided design software (SolidWorks). The model was converted to a stereolithography file and printed by a stereolithography 3D printer (nanoArch P140, BMF Material Technology Inc., Shenzhen, China). Once printed, the nozzles were rinsed with isopropyl alcohol using a syringe to flush the channels. The nozzles were then dried under a stream of air and stored in the dust-free environment.

**8. VWMFs fabrication**

The printable silicone ink was prepared as mentioned above. The syringe with high throughput nozzle was mounted on a custom-built 3D printing setup. The high throughput nozzle was constrained to horizontal movement and maintained the rotational movement of a coated rod which was coated with a layer of release agent to control adjacent hollow filament deposition in the precise location required for fusion. The silicone ink was dispensed from the syringe by means of air pressure generated by an air dispenser (Ultimus V, Nordson EFD) with 45 psi. After printing, the rod was incubated under 80°C for 2 hours and the printed VWMF can be easily released after thermally curing.

**9. Rheology measurement**

A stress-controlled rheometer (ARG2, TA instruments) equipped with cone and plate geometry (50 mm diameter) with an angle β =1 ° is used for rheological studies. The gap at the truncated tip was set at 0.050 mm for all tests. Oscillatory shear rheometry tests are performed at a frequency of 1 Hz.

**10. Mechanical testing**

A tensile test was performed to determine the reliability of VWMFs using a DMA Q800 (TA Znstruments, USA) in the controlled mode for the stress/strain test, in which a ramp force with a slope of 0.5 N/min was applied to the sample. The VWMFs were cut into the pieces with 20 mm length and 4 mm width, and they were flattened between clamps before testing. The stress and strain were determined from the load data and displacement, respectively, by normalizing to the sample length and the cross-sectional area.

**11. Edu (5-ethynyl-2'-deoxyuridine) proliferation assay**

Cells were first cultured on glass or VWMFs for two and five days. Before EdU test, a 2X EdU stock solution was diluted to 10 uM with fresh medium. The samples were next incubated with EdU solution for 2 hours and then endured a standard fixation process. For EdU detection, samples were treated with 1X Click-iT EdU (Yeason, Shanghai, China) buffer at room temperature for 30 minutes and protected from light. After removing the reaction cocktail, the samples were washed with 1 mL of 3% BSA in PBS. The samples were stained with Hoechst and ready for imaging eventually.

**12. Cell culture on VWMFs**

The VWMFs were sterilized using ethanol and then exposed to UV ozone for 7 min. The VWMFs were next immersed in fibronectin solution (50 μg/ mL, Sigma) for 1 hr. Excess fibronectin was washed away by DI water, and the VWMFs were dried with a stream of $N_2$ gas. The fibronectin-coated VWMFs were next transferred to a 12-well plate and immersed in cell culture medium with a high cell concentration up to 0.5 million cells/ml. To achieve an entire 3D cell adhesion, the VWMFs were rotated 120° per hour 3 times in total. The undesired cell attachments on the inner wall of the VWMFs were scratched via a tip before transferring VWMFs to a new 12-well plate. The VWMFs were next immersed into the fresh cell culture medium for a long-term culture and exchange medium every two days.

**13. RNA isolation and quantitative real-time PCR (qRT-PCR) analysis**

Total RNA was isolated from cells using TRIzol (Invitrogen). Quantitative real-time PCR (qRT-PCR) was performed with SYBR Green PCR mastermix. Human GAPDH primers were used as an endogenous control for relative quantifications. Samples in which no expression was detected were given an arbitrary Ct value of 40. All analyses were performed with at least three biological replicates and three technical replicates. Relative expression levels were determined by calculating $2^{-\Delta\Delta Ct}$ with the corresponding s.e.m.

**14. Wound healing assay**

vSMCs were seeded on VMWFs and 35 mm Petri dishes (Corning) for 7 days until approaching 100% confluency. The vertical scratches were made in the confluent monolayer of cells using a glass microcapillary pulled using P2000 glass puller.

**15. OGD (Oxygen Glucose Deprive) assay**

The vSMCs were first incubated under normal conditions for 72 h to achieve 80-90% confluence. Before OGD, to better track the cellular behaviors, vSMCs were labeled with Cell-Tracker Green (Yeason, Shanghai, China). For the OGD groups, the culture dishes were kept at 37 ℃ in an anaerobic and humidified chamber containing a gas mixture of 5% $CO_2$, and 95% $N_2$. The cells were subject to a 4-hour incubation in a

DMEM/F12 medium without glucose. At the end of 4-hour hypoxia exposure, reoxygenation was carried out for 24 h under normal conditions. Meanwhile, the control group was kept in a DMEM/F12 Complete Culture Medium with normal oxygen level. Time-lapse video microscopy was performed by GE Ultra Live-cell Imaging System.

**16. Scanning Electron Microscope (SEM)**

Structure and microscopic features of the fibers were investigated by field emission scanning electron microscopy (FE-SEM) (FEITeneo, FEICo., Hillsboro, OR and Carl Zeiss Microscopy GmbH, Germany). Fully dried samples were sputter-coated with a ~10 nm thick gold layer, to ensure conductivity.

**17. Image Analysis**

Phase and fluorescence images were recorded using an inverted epifluorescence microscope (Leica DMi8; Leica Microsystems) equipped with a monochrome charge-coupled device (CCD) camera. ImageJ (NIH, Bethesda, MD) was used for the measurement of cell areas.

**18. Statistics**

Statistical analysis was performed using GraphPad Prism. For statistical comparations between two data sets, P-values were calculated using the student t-test function. Whereas for three or more data sets, P-values were calculated using the one-way ANOVA with Tukey post-hoc analysis.
